# The TyphiNET data visualisation dashboard: Unlocking *Salmonella* Typhi genomics data to support public health

**DOI:** 10.1101/2024.06.03.595798

**Authors:** Zoe A. Dyson, Louise Cerdeira, Vandana Sharma, Megan E. Carey, Kathryn E. Holt, Global Typhoid Genomics Consortium

**Author notes:** Correspondence Zoe A. Dyson & Kathryn E. Holt. Indicates equal contribution (Zoe A. Dyson, Louise Cerdeira).

## Abstract

**Background:** *Salmonella enterica* subspecies *enterica* serovar Typhi (abbreviated as ‘Typhi’) is the bacterial agent of typhoid fever. Effective antimicrobial therapy reduces complications and mortality; however, antimicrobial resistance (AMR) is a major problem in many endemic countries. Prevention through vaccination is possible through recently-licensed Gavi-supported typhoid conjugate vaccines (TCVs), and national immunisation programs are currently being considered or deployed in several countries where AMR prevalence is known to be high. Pathogen whole genome sequence data are a rich source of information on Typhi variants (genotypes or lineages), AMR prevalence, and mechanisms. However, this information is currently not readily accessible to non-genomics experts, including those driving vaccine implementation or empirical therapy guidance.

**Results:** We developed TyphiNET (https://www.typhi.net), an interactive online dashboard for exploring Typhi genotype and AMR distributions derived from publicly available pathogen genome sequences. TyphiNET allows users to explore country-level summaries such as the frequency of pathogen lineages, temporal trends in resistance to clinically relevant antimicrobials, and the specific variants and mechanisms underlying emergent AMR trends. User-driven plots and session reports can be downloaded for ease of sharing. Importantly, TyphiNET is populated by high-quality genome data curated by the Global Typhoid Pathogen Genomics Consortium, analysed using the Pathogenwatch platform, and identified as coming from non-targeted sampling frames that are suitable for estimating AMR prevalence amongst Typhi infections (no personal data is included in the platform). As of February 2024, data from a total of n=11,836 genomes from 101 countries are available in TyphiNET. We outline case studies illustrating how the dashboard can be used to explore these data and gain insights of relevance to both researchers and public health policy-makers.

**Conclusions:** The TyphiNET dashboard provides an interactive platform for accessing genome-derived data on pathogen variant frequencies to inform typhoid control and intervention strategies. The platform is extensible in terms of both data and features, and provides a model for making complex bacterial genome-derived data accessible to a wide audience.

## Background

*Salmonella enterica* subspecies *enterica* serovar Typhi (abbreviated as ‘Typhi’) is the bacterial agent of typhoid fever [1], a faeco-orally transmitted systemic bacterial infection, which sickens an estimated nine million people each year [2]. Most illnesses occur in low- to middle-income countries (LMIC) in settings with insufficient sanitation infrastructure, microbiologically unsafe water and food, and poor hygiene, where the disease burden is highest among children [3]. Effective antimicrobial therapy hastens the resolution of typhoid symptoms [4–6], reduces the risk of complications [7], and reduces mortality from ∼10% to 1% [8,9]. Formal diagnosis and susceptibility testing requires blood culture that has low sensitivity (<60%) and is often limited or unavailable in high-burden settings [10,11].

Therefore, therapy is often empiric, and guided by local antimicrobial resistance (AMR) patterns rather than direct testing. For example, the World Health Organization (WHO) ‘AWaRe (Access, Watch, Reserve) Antibiotic Book’ [12] recommends treating suspected typhoid with ciprofloxacin if the local prevalence of resistance is low, and oral azithromycin for uncomplicated disease or intravenous ceftriaxone for severe disease if local prevalence of ciprofloxacin resistance is high. The former first-line drugs ampicillin, chloramphenicol, and trimethoprim-sulfamethoxazole have not been recommended by the WHO for typhoid fever since the 1990s, when multidrug resistance (MDR, defined as resistance to these three agents) became common [5,13]. Extensively drug resistant (XDR) strains have been reported and these are MDR strains that are resistant to ciprofloxacin and ceftriaxone. Patients with uncomplicated disease due to XDR Typhi may be treated with azithromycin and carbapenems are used for severe disease but are problematic due to cost and the need for intravenous administration.

While vaccines to prevent typhoid have been available for decades, they have not been widely implemented in endemic regions. The situation is changing now due to the prequalification of typhoid conjugate vaccines (TCVs) by the World Health Organization (WHO) in 2018. TCVs are safe and effective in children and in infants as young as six months of age [13]. TCVs are now eligible for Gavi support, providing a potentially affordable route for low-income countries to invest in typhoid prevention through national immunisation programs. As AMR has an impact on the clinical outcomes of Typhi infections, it is not only the burden of typhoid fever but also the prevalence of AMR that needs to be considered when weighing the costs and benefits of disease prevention through vaccination [14].

Indeed, TCV was the first vaccine to be recommended by the WHO based partially on pathogen-specific AMR concerns. For example, XDR typhoid outbreaks in Pakistan and ciprofloxacin resistant (CipR) outbreaks in Zimbabwe prompted responsive TCV roll-out in affected areas, which were effective in reducing local disease incidence [15–20]. Such responses have since been followed by introduction of national immunisation programs, with Pakistan being the first country to introduce TCV into its routine immunisation schedule.

Country-level AMR prevalence data are important to inform both empiric treatment of typhoid fever, and make the case for investment in national immunisation programs. *Salmonella enterica* is included in the WHO Global Antimicrobial Resistance and Use Surveillance System (GLASS) [21], but is not currently disaggregated by serovar. Local data on Typhi AMR remain scarce in most LMICs, and are mostly gathered in the context of time-limited research studies, outbreak investigations, or from travellers returning to other countries that have routine surveillance [22]. While informative, these data are not collected consistently, are predominantly based on phenotypic testing, and methods and interpretive criteria can vary by country and over time, which complicates reporting and interpretation. Whole genome sequencing (WGS) data are increasingly adopted as the standard for strain characterisation of Typhi [23–25], and a hierarchical genotyping and nomenclature scheme (GenoTyphi) has been developed to aid the detection and tracking of lineage variants [26,27]. The genetic determinants of AMR in Typhi are well understood [5,23,28], such that AMR phenotypes can also be predicted from WGS data, with a recent study demonstrating 99.9% concordance between AMR genotypes and phenotypes assessed in the English reference laboratory [25]. Consequently, WGS is now a standard method of characterisation in routine surveillance of typhoid in reference laboratories in many high-income countries [22,25,29,30], as well as in research studies globally, making WGS data a rich source of information on Typhi pathogen diversity and local AMR patterns [23].

As AMR continues to evolve and spread, WGS-based pathogen surveillance has the potential to inform public health policies for typhoid such as empirical therapy guidelines [31]; water, sanitation, and hygiene (WASH) interventions that could impact pathogen transmission [32]; and national vaccine introduction decision making [33,34]. However, at present, these data are not universally accessible to decision-makers at different levels of public health policy nor presented in a format that can inform on national prevalence of AMR. Typhi genome data are browsable in various public databases including the National Center for Biotechnology Information (NCBI) Pathogen Detection portal [35], Enterobase [36,37], BIGSdb [38], and Pathogenwatch [28]. However, these databases (i) are designed for a user base with expertise in genomics, bioinformatics, and WGS analysis; (ii) include WGS data from heterogeneous sources, including many that are not suitable for surveillance of AMR prevalence (e.g., outbreak investigations [39], or studies that specifically sequence resistant strains to ascertain mechanisms [40]); and (iii) do not provide country-level summaries of key variables such as genotype and AMR prevalences since uncurated bundles of input data are not suitable for this.

Here, we present the TyphiNET genomic surveillance dashboard for typhoid (available at: https://www.typhi.net), which aims to make genome-derived data on Typhi genotypes and AMR accessible to a broad user-base. TyphiNET is powered by existing informatics solutions for Typhi genomic analysis (including the GenoTyphi framework and Pathogenwatch platform), and leverages contextual metadata collected and curated by the Global Typhoid Genomics Consortium (GTGC), to filter and analyse raw heterogeneous public genome data and extract meaningful AMR and genotype (variant) prevalence. Where sufficient input data are available, the dashboard also allows visualisation of temporal trends, and users can interrogate the association of specific AMR determinants with genotype backgrounds.

## Implementation

### Dashboard architecture

TyphiNET was developed as an open-source MERN (MongoDB, Express, React, Node) stack JavaScript application (**Fig 1**). Front-end visualisations are implemented via ReactJS libraries, while back-end operations are implemented using ExpressJS and NodeJS. Genome-derived AMR and genotype data, and curated contextual metadata (see below), are retrieved from Pathogenwatch (https://pathogen.watch) via an automatic robot for web scraping, dubbed Spyder v2.0 (https://github.com/lcerdeira/Spyder) [41], and injected into the TyphiNET MongoDB Atlas via the back-end. The web application is deployed using the Heroku platform. All code is freely available under a GNU-GPL 3.0 licence at https://github.com/typhoidgenomics/TyphiNET. The version described in this manuscript is v1.5.1, DOI: 10.5281/zenodo.10667321, which includes data updated on February 15th 2024 [42].

**Figure 1.**
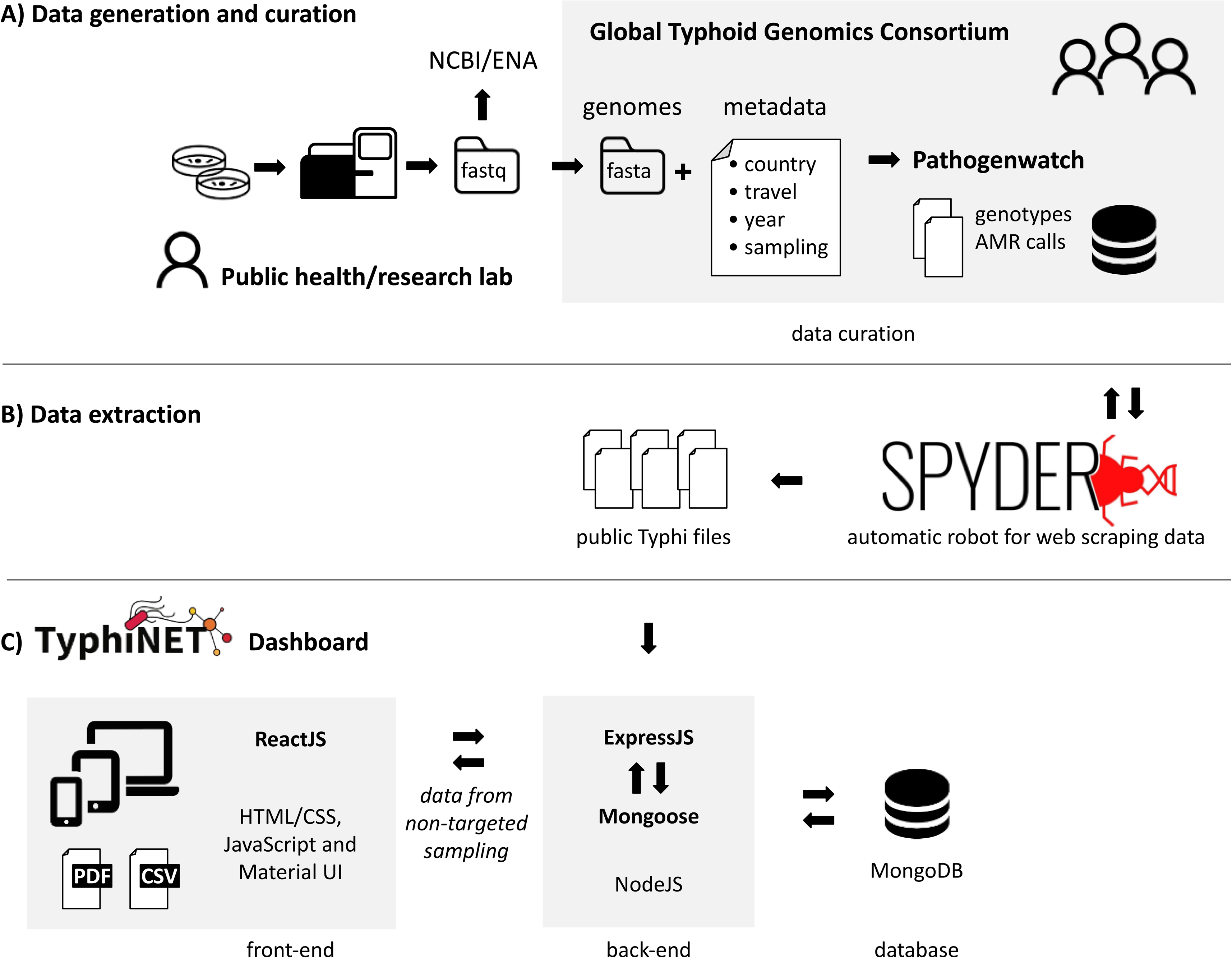
TyphiNET data curation and dashboard architecture. **(A)** The Global Typhoid Genomics Consortium (GTGC) aggregates and curates Typhi genome data and metadata, using Pathogenwatch as both an analysis platform (calling genotypes and AMR determinants from genome assemblies) and publicly accessible data store. Metadata that are not typically available in NCBI/ENA but are collected and curated by the GTGC include purpose of sampling (to tag datasets that are suitable for estimating AMR/genotype prevalence) and information on country-of-travel for travel-associated isolates (to identify country of origin). **(B)** A web-scraper is used to pull the latest versions of genotypes, AMR determinants, and metadata files from GTGC-curated Typhi collections in Pathogenwatch, which are used to populate the TyphiNET database. **(C)** The TyphiNET dashboard is implemented as a MERN (MongoDB, Express, React, Node) stack JavaScript application as illustrated. Genome data are filtered to exclude low-quality genome sequences, and data sets whose sampling frames make them unsuitable for AMR surveillance (such as those targeted towards sequencing of resistant strains only, or outbreak investigations), before calculating national/annual prevalences of AMR and genotypes to display in interactive plots. ReactJS is used to provide user interface layouts suitable for viewing the interactive plots on a range of devices (computer, tablet, phone). Users can also download static images of current plot displays (PNG), static reports with all current plots (PDF format), or a copy of the TyphiNET database (CSV format).

### Data curation and processing

The GTGC aggregates Typhi WGS data to facilitate monitoring the emergence and spread of AMR and inform targeted public health action against typhoid fever [23]. WGS data generated by research or public health laboratories and deposited in International Nucleotide Sequence Database Collaboration (INSDC) databases (i.e., NCBI, EMBL-EBI, DDBJ) are assembled and quality filtered by the GTGC [23], and uploaded to Pathogenwatch (https://pathogen.watch) for analysis (**Fig 1**). Typhi Pathogenwatch screens assemblies for known determinants of AMR [28] and carries out lineage assignment according to the GenoTyphi genotyping framework [27,43].

The GTGC also curates contextual metadata (i.e., source data) associated with each Typhi genome sequence, to enhance the re-usability of the WGS data. This is done via requesting data generators (most of whom are GTGC members) to complete a standardised metadata template (available at https://bit.ly/typhiMeta). Key fields in the GTGC metadata template that are not commonly or consistently included in metadata submitted to INSDC or supporting publications, but which are important for re-using genome data for AMR surveillance, are (i) travel information (Travel Associated: yes or no, Country of Travel); (ii) purpose of sampling (Targeted: Cluster Investigation, AMR investigation, Other; or Non Targeted: Reference lab, Surveillance Study, Routine diagnostics, Other); (iii) identifying repeat isolates; and (iv) data to confirm case status (Host Health State: Symptomatic or Asymptomatic Carrier; Source: Blood, Stool, Environment or Food). Repeat isolates, defined as those that represent the same occurrence of typhoid infection, are excluded such that only a single ’primary’ isolate (either the first, or the best quality genome, for each unique case) are included in the GTGC data set [23]. ‘Country of origin’ is defined as the country where the pathogen was isolated, or for travel-associated infections, the country recorded as the presumed country of infection based on travel history [22,44–46]. The GTGC-curated metadata (including ‘Country of Origin’, and with repeat isolates removed) is processed and uploaded to Pathogenwatch along with the sequence data, to generate curated ‘Collections’ in Pathogenwatch (one for each source study or public health lab). The Spyder API [41] is then used to retrieve data files from Pathogenwatch for the GTGC-curated collections, for injection into the TyphiNET database.

In the TyphiNET dashboard, genomes are further filtered to include only those recorded as coming from non-targeted sampling frames (Reference lab, Surveillance Study, Routine diagnostics, Other). Those from targeted sampling frames (Cluster investigation, AMR focused, Other) or unspecified sampling frames are excluded from dashboard analyses and visualisations (although they are included in the database download, for completeness). The TyphiNET dashboard also filters out cases recorded as asymptomatic carriers (n=119) or coming from gallbladder (n=1) or environmental (n=14) samples; the rest are assumed to represent acute illness (including n=9,039 recorded explicitly as blood isolates and/or symptomatic typhoid).

### Statistical and design considerations

Typhoid intervention strategies such as immunisation programs and changes in empirical therapy are typically implemented at a country level, by national immunisation technical advisory groups (NITAGs) and ministries of health [47]. The TyphiNET dashboard therefore focuses on country as the geographical unit, to report national annual prevalences of genotypes and AMR aggregated from all available data sources for that country. Prevalence estimates are simple proportions (expressed as percentages), calculated from the data in the curated TyphiNET database (i.e. non-repeat assumed-acute cases from non-targeted sampling frames, as outlined above). Where multiple data sources are available for a given country, the prevalence is a simple weighted pooled estimate calculated by summing the numerators and denominators across all available data for the given country and the selected time period. The minimum sample size to report a national annual prevalence is N≥10 (equivalent to current WHO GLASS reports [21], which require data from 10 individuals in order to report a national prevalence rate). The minimum sample size to report a national prevalence on the map is N≥20. The number of samples, and number and scope of available data sources, varies substantially by country; however, previous robustness analyses reported by the GTGC show that, for countries with multiple data sources (e.g., burden studies in different cities, and returning traveller data collected in other countries), the per-data-set prevalence estimates are largely concordant with the pooled country-wide estimates [23].

### AMR definitions

Binary variables representing predicted resistance phenotypes are calculated from the AMR genes and Single Nucleotide Polymorphisms (SNPs) reported by Pathogenwatch [28], as summarised in **Table 1**. Note that there is limited data to assess the clinical significance of *acrB* mutations on treatment response to azithromycin, for simplicity we refer to isolates carrying these as azithromycin resistant [40].

**Table 1.**
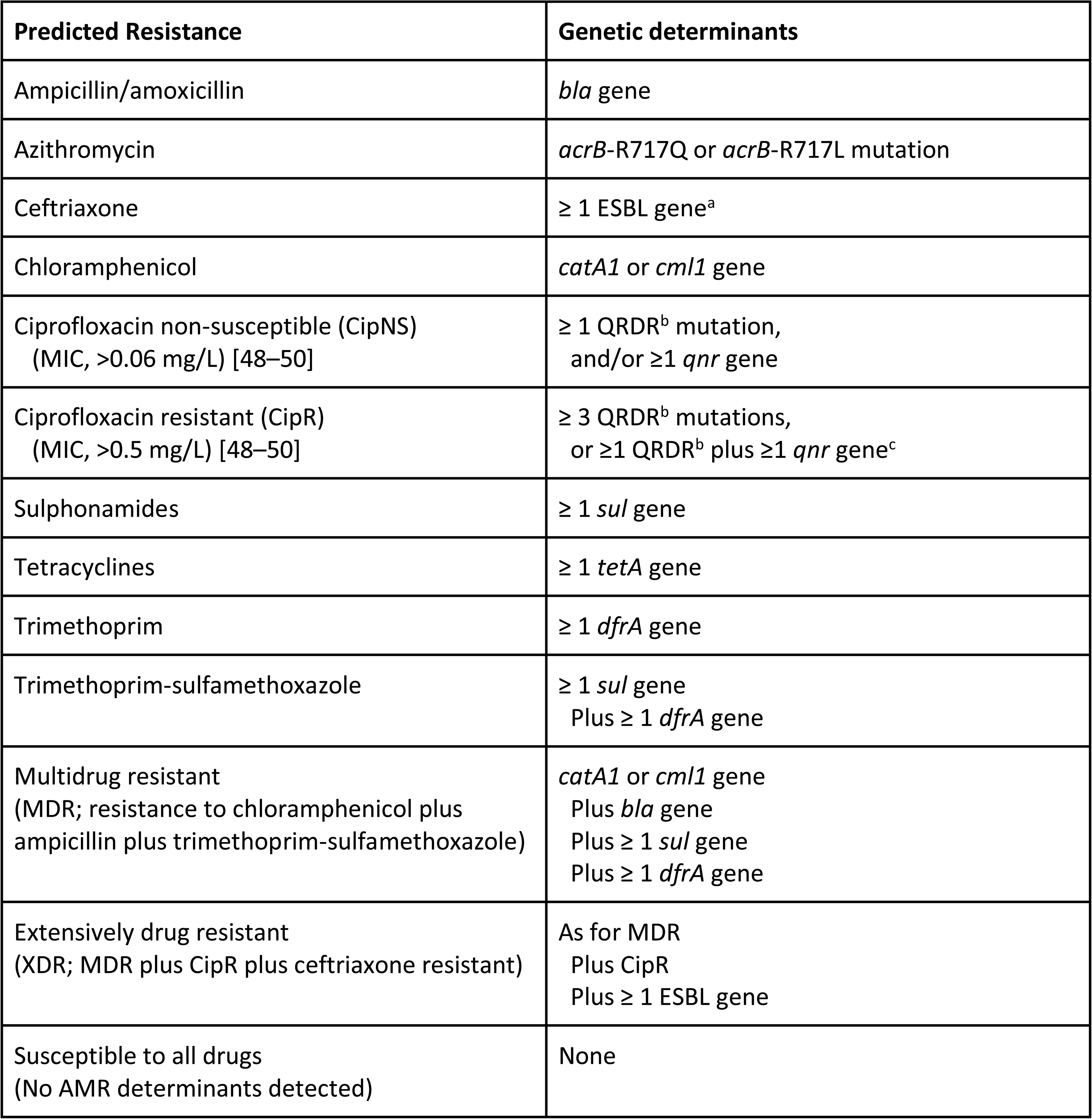
Definitions used to calculate AMR variables. Genetic determinants reported by Pathogenwatch are used to calculate binary resistance prediction variables in TyphiNET. ^a^ESBL = extended spectrum beta-lactamase gene (those currently detected in the Typhi genomes are *bla*_CTX-M-12_*, bla*_CTX-M-15_*, bla*_CTX-M-55_*, bla*_OXA-7_*, bla*_OXA-134_*, bla*_SHV-12_). ^b^QRDR = quinolone resistance determining region mutations tracked by Pathogenwatch [28] (these are currently *gyrA* S83 and D87, *gyrB* S464, *parC* S80 and E84).

### Global map view

The first visualisation panel summarises country-level data on a world map. Users select from a dropdown menu which variable to colour countries by: (i) national prevalences of clinically relevant AMR profiles including MDR, XDR, CipNS, CipR, AziR, susceptible to all drugs (see **Table 1**), (ii) national prevalence of genotypes (lineage variants), including the dominant genotype per country and the prevalence of genotype 4.3.1; or (iii) number of samples available. By default, the map view shows values enumerated from all Typhi isolates, sampled both locally in-country and travel-associated cases captured in other countries. Users can choose to filter to either local or travel data only (via a toggle button) and/or to filter on a specified time window (by selecting start and end years). These filters apply to all dashboard plots, not just the map.

AMR prevalences per country are indicated visually on the world map, using increasing colour intensity to signal categorical prevalence ranges of escalating concern with respect to use of the drug for empiric therapy: (i) 0, no resistance detected; (ii) >0 and ≤2%, resistance present but rare; (iii) >2 and ≤10%, resistance uncommon; (iv) >10% and ≤50%, resistance common; (v) >50%, established resistance. The ‘sensitive to all drugs’ plot (selected via the ‘map view’ dropdown menu) is coloured differently, to draw attention to countries with low prevalence of pan-susceptible strains and thus where choice of antimicrobial is most important: (i) <10% pan-susceptible; (ii) >10 and ≤20%; (iii) >20 and ≤50%; (iv) >50 and ≤90%; >90%. Prevalence estimates are visualised where ≥20 sequences are available for a given country and timeframe, otherwise “Insufficient data” is shown (light grey; **Fig S1a**).

Users can interact with the map by hovering the mouse cursor over a country, to view a tool-tip displaying the name of the country and the number (N) and percentage of genomes from that country that are resistant (or pan-susceptible or H58/4.3.1, depending on the ‘map view’ variable selected); numbers shown always reflect the current choice of local/travel and temporal filters. Selecting ‘No. Samples’ as the ‘map view’ variable to plot colours the map according to number of samples available using the current filters; in this view, hovering over a country reveals a tool-tip displaying the name of the country, number of samples and genotypes, and prevalence of H58/4.3.1 and each of the AMR categories.

### Detailed plots (for country-level data)

The second visualisation panel includes four additional data plots designed to highlight annual trends in genotype and AMR prevalences, AMR prevalence within genotype, and molecular mechanisms underlying AMR. Upon loading the dashboard, these plots are populated by the full dataset (i.e., all countries), however, they were designed mainly for the purpose of showing detail for a single country of interest. Users can select a country by clicking it on the map, or selecting its name from the ‘select country’ dropdown menu below the map. The data plots are then populated by filtering the database to include only samples from the selected country of origin (along with applying any local/travel and time filters selected in the map panel).

The ‘Drug resistance trends’ plot (**Fig S1c**) shows annual pooled global prevalence of genomically-predicted resistance to the drugs listed in **Table 1** (as well as prevalence of genomes identified as ‘susceptible to all drugs’). By default, trend lines are plotted for the most currently relevant AMR categories (MDR, XDR, CipR, CipNS, AziR, CefR, Trimethoprim-sulfamethoxazole resistant, Susceptible), but individual variables can be hidden or displayed via the dropdown menu ‘Select drugs/classes to display’. For the remaining three plots (**Fig S1b,d,e**), users can visualise data as either counts or percentages using dropdown menus.

For the ‘Resistance frequencies within genotypes’ and ‘Resistance determinants within genotypes’ plots, percentages are shown by default to highlight the most resistant pathogen variants. By default, the ‘Resistance frequencies within genotypes’ plot shows the top five most-resistant genotypes in the currently-selected country, but other genotypes can be selected via ‘data view’ dropdown menu. The ‘Resistance determinants within genotypes’ plot shows the genetic determinants underlying resistance in up to the 10 most-resistant genotypes for the currently-selected filters and a selected drug category. The default view is of determinants conferring non-susceptibility to ciprofloxacin, as this is the most relevant to empiric treatment choice, however other drugs can be selected from the ‘drug class’ dropdown menu. For the ‘Genotype distributions’ plot, counts are shown by default to mimic epidemic curves used in epidemiological investigations for case counts. As for the map views, hovering the mouse cursor over these plots reveals a tool-tip displaying the raw data count (N) and prevalence (%) underlying each data point.

### Static outputs

User-generated visualisations can be downloaded individually as portable network graphics (PNG) files (**Fig S1f**), and a report of all current data visualisations can be downloaded as a portable documents format (PDF) file (example output in **Data S1**). If a country is selected, the report includes a list of publications (PubMed IDs) for genomes included in the current view, to facilitate proper citation and provenance-tracking of constituent datasets (otherwise the report refers readers to download the database for this information). A plain text line list (comma separated values, CSV) of the full TyphiNET database is also available for download (example output in **Table S1**). This file contains the GTGC-curated sample metadata (including country of origin, year of isolation, purpose of sampling, travel association; provenance information for individual genomes such as originating lab, primary publication PubMed ID, sequence data accession numbers), as well as genome-derived AMR determinants and pathogen genotypes assigned by Pathogenwatch. This file is intended to facilitate provenance-tracking of individual genome sequences, and to allow expert users to further explore the data using other tools.

## Results and discussion

Version 1.5.1 of the TyphiNET database (February 2024) included data derived from 12,671 genomes curated by the GTGC, including those described in the initial GTGC publication [23] and more recent published datasets from Kenya [51] and Fiji [52]. The TyphiNET dashboard displays data from genomes from assumed-acute typhoid cases from non-targeted sampling frames (see **Implementation**), resulting in a total of n=11,836 Typhi in version v1.5.1. The filtered database contains data from 101 countries, however as the map view of sample counts shows (**Fig 2a**), most countries are represented by very few samples. Prevalence estimates are calculated only for countries represented by ≥20 sequences, currently n=30 countries (see e.g., map of XDR prevalence, **Fig 2b**). The database currently includes samples from 1958-2021, with the majority from 2010 onwards (n=10,382, 87.7%). High-burden countries in South Asia are well represented with >1,300 genomes each (Bangladesh, n=1,664; India, n=2,327; Nepal, n=1,300; Pakistan, n=1,526), including both local data from large-scale disease-burden studies [53,54] and travel-associated infection isolates sequenced in other countries (Bangladesh, 14.1%; India, 45.2%; Nepal, 1.8%; Pakistan, 39.6%) [22,25]. African countries are currently represented by low numbers of genomes, with ten countries exceeding 20 genomes (Malawi n= 568; Kenya n=824; Nigeria n=170; Ghana n=69; Rwanda n=52; South Africa n=312; Uganda n=36; Tanzania n=33; Cameroon n=27; Gambia n=24). This highlights the need for culture-based surveillance studies in typhoid endemic countries in Africa [55]. Notably TyphiNET and the GTGC provide a mechanism for continued updating of the database as new genomes are released from such efforts, as well as from travel-associated infections captured in other countries. For example, Nigeria is currently represented by n=28 travel-associated infections and n=142 local infections, which reflect the same general trends in terms of dominance by genotypes 3.1.1 and 2.3.1, with higher prevalence of MDR in 3.1.1 highlighted by both data sources (**Fig S2**).

**Figure 2.**
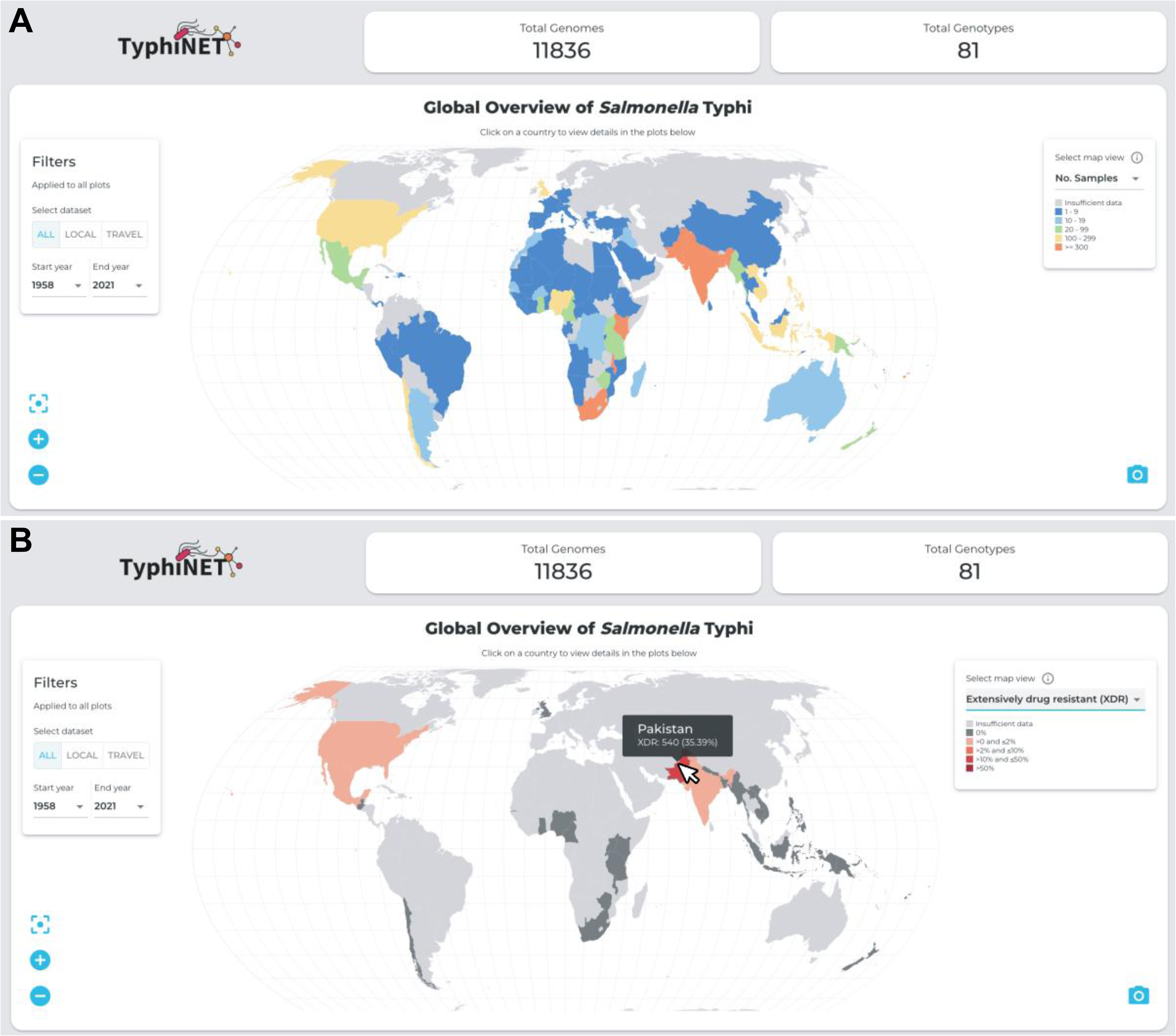
Global views of data gaps and AMR prevalence. **(A) Total sample counts per country.** Top panels indicate the number of sequences and genotypes present in the TyphiNET dashboard as of February 2024. Left panel indicates controls for filtering the data visualised by data source (all data, locally collected cases, or travel-associated cases) and time period (by providing start and end years for the period). Countries on the map are coloured by the total number of samples as per the inset legend (top right of map). **(B) National frequencies of XDR.** Countries on the map are coloured by XDR frequency as per the inset legend (top right of map). Data are shown where there are ≥20 sequences available for the country of interest. Tool tip indicates summary statistics for Pakistan upon mouse over.

We conducted an informal assessment of dashboard useability by asking untrained users from different geographies to explore the dashboard and use it to answer 10 multiple-choice questions. Questions were designed to assess whether users could successfully interact with the dashboard in order to find the answers to specific questions regarding the prevalence, trends, and determinants of typhoid fever resistance in specific countries and time periods (**Table 2**). Our goal was to assess the ability of users who are generally familiar with typhoid and AMR to find the specific information they need, rather than to assess comprehension or understanding of typhoid and AMR. Therefore the audience for the quiz was members of the Global Typhoid Genomics Consortium and their colleagues at academic and public health institutions, and we did not track the professional background or expertise of individual respondents, nor did we provide any training or background to typhoid fever or the concepts employed in the dashboard. The results (from n=42 respondents) suggest the dashboard is sufficiently intuitive for users who are familiar with the concepts, but not familiar with the dashboard interface, to find the correct answers to these types of questions (>90% correct responses to 8/10 questions, see **Table 2**).

**Table 2.** Informal assessment of dashboard useability. ^1^This question has no objective answer, as the prevalence of genotype 3.1.1 in Nigeria moves up and down between 2010-2020.

| Question | Answers (*Correct) | Correct responses N (%) |
| --- | --- | --- |
| In which country is extensively drug-resistant (XDR) *S. Typhi* most prevalent (across all years)? | India<br>Mexico<br>Pakistan *<br>United States | 41 (98%) |
| How many total genomes are available from Nepal since 2010? | 1100<br>1274 *<br>1484<br>1660 | 39 (93%) |
| How many of the genomes available from Nepal since 2010 are from travel data? | 20 *<br>45<br>75<br>103 | 41 (98%) |
| What percentage of *S. Typhi* from Kenya isolated between 2010 and 2020 were multidrug resistant (MDR)? | 23.45%<br>43.29%<br>78.41% *<br>100% | 39 (93%) |
| What is the overall trend in Ciprofloxacin non-susceptibility (CipNS) in Kenya from 2010 to 2020? | Increasing *<br>Decreasing<br>No change | 41 (98%) |
| Which genotypes had the highest rates of Ciprofloxacin non-susceptibility (CipNS) in Kenya from 2010-2020? (two answers) | 4.3.1.1.EA1<br>4.3.1.2.EA2 *<br>4.3.1.2.EA3 *<br>0.0.1<br>4.3.1.1.EA2<br>4.3.1.1.EA3 | 34 (81%) selected ONE or BOTH correct genotypes |
| What was the most common genotype in Nigeria in 2013? | 2.3.1<br>2.3.2<br>3.1.1 *<br>4.1 | 41 (98%) |
| <sup>1</sup>What is the overall trend in prevalence of this genotype in Nigeria between 2010 and 2020? | Increasing<br>Decreasing<br>Stable | 11 (26%)<br>17 (16%)<br>24 (57%) |
| What is the prevalence of trimethoprim resistance in this genotype in Nigeria between 2010 and 2020? | 45%<br>78%<br>89% * | 32 (76%) |
|  | 100% |  |
| Which resistance gene is responsible for the majority of trimethoprim resistance in this genotype in Nigeria between 2010 and 2020? | dfrA14 + sul2<br>dfrA15 *<br>dfrA1 + sul1<br>dfrA1 + sul1 + sul2 | 40 (95%) |

In the following sections, we present three case studies that highlight the utility of the TyphiNET dashboard as a tool for understanding national AMR trends and the underlying mechanisms and pathogen genotypes. The case studies were developed to illustrate previously documented shifts in local Typhi populations in African and Asian countries, each associated with different types of changes in AMR patterns (emergence or decline of resistance, and lineage replacement) that have implications for local typhoid control.

### Case study 1: Emergence and clonal expansion of XDR typhoid in Pakistan

Selecting ‘Extensively drug resistant’ from the dropdown menu in the map, it is clear that Pakistan has many XDR cases (**Fig 2b**). Toggling the ‘Local’ and ‘Travel’ filters shows that this high prevalence is evident in data from both sources. Clicking on Pakistan in the world map allows exploration of the genotypes, resistance mechanisms and annual trends underlying the high prevalence of XDR in the country. With the filter set to ‘All’ (i.e., including local and travel data) and year range from 2010 to 2021, the ‘Resistance trends’ plot shows there is sufficient data (N≥10) per year from 2014-2020 to calculate annual prevalence values. The plot shows the first emergence of XDR in Pakistan in 2016 (**Fig S3a**), against a background of persistently high CipNS prevalence (>96% throughout 2014-2020) and MDR (∼60% in 2014- 2017, rising to 84% in 2020). The ‘Resistance frequencies within genotypes’ plot shows MDR focused in the 4.3.1.1 genotype background (**Fig 3a, Fig S3b**), with CefR, CipR and XDR (ie the combination of MDR+CefR+CipR) localised in the derived genotype 4.3.1.1.P1. The ‘Genotype distribution’ plot, with data view set to ‘Percentage per year’, shows the first appearance of genotype 4.3.1.1.P1 in 2016 (**Fig 3b, Fig S3c**). Mousing over the year 2016 brings up the tooltip, which clarifies that there is a single genome of 4.3.1.1.P1 amongst total N=96 for the year. The plot shows prevalence of 4.3.1.1.P1 increased in subsequent years, reaching 87.5% in 2020, with the minor genotypes reducing in prevalence continuing to persist.

**Figure 3.**
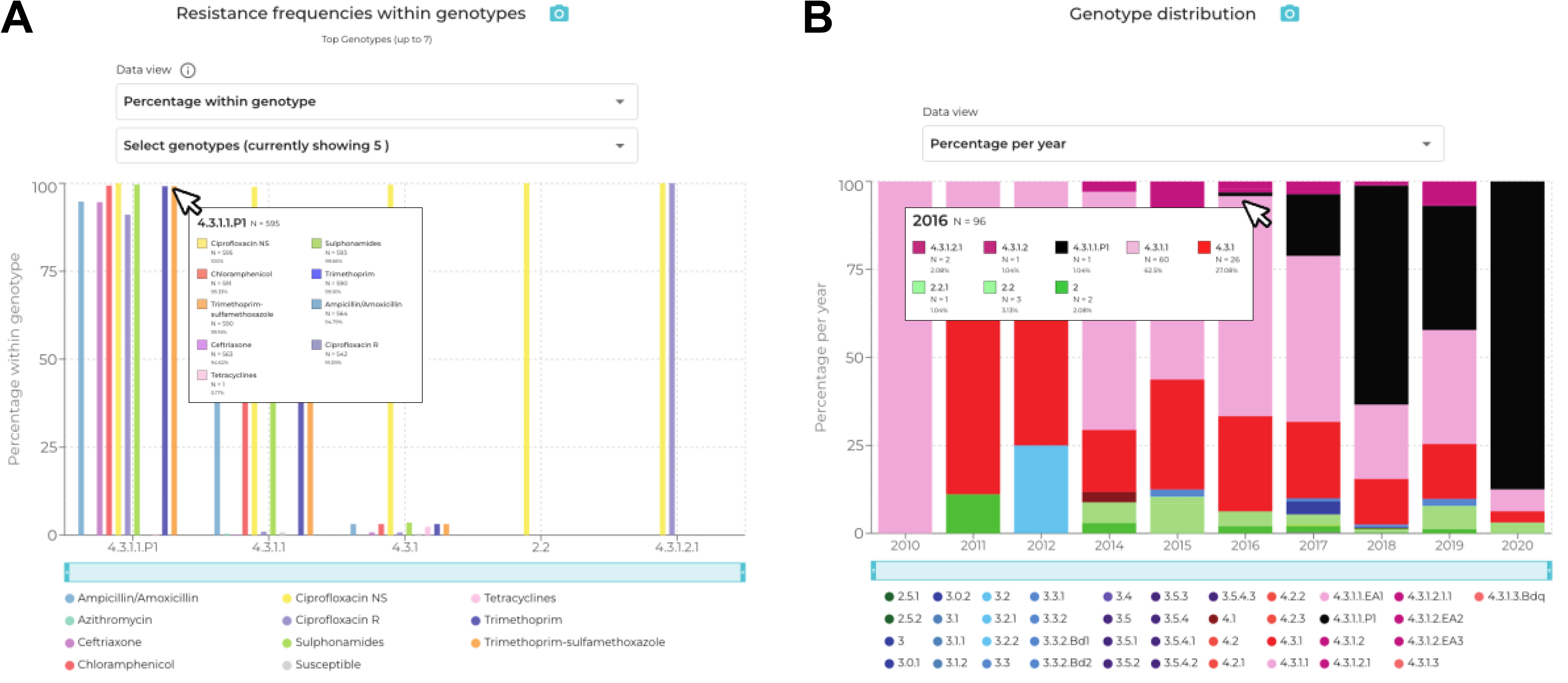
Exploring the emergence of XDR Typhi in Pakistan with the TyphiNET dashboard. **(A) ‘Resistance frequencies within genotypes’ plot shows frequencies of resistance to different drug classes, within common genotypes circulating in Pakistan.** Bars are coloured according to the inset legend. **(B) ‘Annual genotype distribution’ plot shows the frequencies of pathogen genotypes circulating in Pakistan per year.** Genotypes are coloured as per the inset legend.

The ‘Resistance determinants within genotypes’ plot shows the genetic basis for XDR in 4.3.1.1.P1. The default view shows fluoroquinolone resistance determinants, and highlights that CipR in the genotype 4.3.1.1.P1 is due to the combination of a single QRDR mutation and a *qnrS* gene (**Fig S3d**). Most other genotypes also have a single QRDR mutation (resulting in CipNS, shown in yellow), indicating that high-level resistance could emerge in any of these strain backgrounds through additional *gyrA*/*parC* mutations [56,57] or plasmid-mediated acquisition of *qnr* genes[39,58,59]. Interestingly, the plot also highlights the presence of a variant with three QRDR mutations (resulting in CipR, shown in red), genotype 4.3.1.2.1, which emerged in India some decades ago [23,60] (the genotype timeline shows this was detected in Pakistan between 2016-2019, in parallel with XDR 4.3.1.1.P1). Selecting ‘Ceftriaxone’ from the dropdown menu shows resistance in 4.3.1.1.P1 is mainly due to *bla*_CTX-M-15_ (**Fig S3e**); the same gene is found in a single isolate of 4.3.1.1 and two of 4.3.1.

Selecting ‘Trimethoprim-sulfamethoxazole’ from the dropdown menu shows resistance in 4.3.1.1.P1 is mostly due to *dfrA7* plus *sul1* and *sul2*, and that this combination is also common in the parent genotype 4.3.1.1 (**Fig S3f**), consistent with emergence of XDR by acquisition of *qnrS*+*bla*_CTX-M-15_ within the locally circulating MDR+QRDR 4.3.1.1 variant.

The TyphiNET dashboard conveys several key points about XDR typhoid, which reflect the emerging picture of the problem captured in the wider literature [39,54,61–63]. These include (i) time of emergence of XDR typhoid; (ii) that XDR cases are due to emergence and dissemination of a single variant, genotype 4.3.1.1.P1; (iii) the underlying mechanisms of resistance; (iv) that this genotype shares with its parent, 4.3.1.1, the MDR genes and single QRDR mutation but has acquired *qnrS* and *bla*_CTX-M-15_ to become XDR; (v) that the XDR 4.3.1.1.P1 has not established locally transmitting populations outside Pakistan, at least not by 2020 (this is still true as of April 2024, however more recent data are sparse due to a decline in travel and prioritisation of SARS-CoV-2 sequencing during the COVID19 pandemic). Intercontinental transmission of XDR typhoid associated with travel has been reported in several countries [22,29,45,64–67], however, to date only a single report of a localised outbreak has occurred outside Pakistan [68]. TyphiNET will provide a means of monitoring the emergence of new XDR strains, as well as the persistence of XDR Typhi 4.3.1.1.P1 within Pakistan and its eventual spread to other settings which may motivate more widespread TCV use. The latter is of critical importance as recent surveillance data have shown that ancestral populations of genotype 4.3.1.1 circulating in Pakistan have acquired *acrB* mutations conferring azithromycin resistance [69] (as can be seen by selecting ‘Azithromycin’ in the ‘Resistance determinants within genotypes’ plot), suggesting that 4.3.1.1.P1 may also be able to tolerate these mutations.

### Case study 2: Decline of MDR and emergence of azithromycin resistance in Bangladesh

Selecting ‘Multidrug resistant’ from the dropdown menu in the map, it can be seen that MDR infections are distributed throughout parts of sub-Saharan Africa, South-eastern Asia and South Asia (**Fig S4a**). Toggling the ‘Local’ and ‘Travel’ filters reveal a high prevalence of MDR cases in Bangladesh from both data sources. Clicking on Bangladesh in the world map, or using the ‘select country’ dropdown menu, allows exploration of country-level trends over time. With the filter set to ‘All’ (i.e., including local and travel data) and year range from 2005 to 2021, the ‘Resistance trends’ plot shows there are sufficient data (N≥10 per year) to calculate annual AMR prevalences from 2005-2019 (with the exception of 2006) (**Fig 4a, S4b**). Mousing over the year 2005 reveals MDR (deep red line) was common in 2005 (91%), declining in subsequent years to 8-30% between 2013 and 2019. At the same time, CipNS (yellow line) remained high (>92%) throughout 2005-2019 (**Fig S4b**). The ‘Genotype distribution’ plot, with data view set to ‘Percentage per year’ shows that 4.3.1 genotypes, including sublineages 4.3.1.1, 4.3.1.2, and 4.3.1.3, have also been gradually declining over the same time period (**Fig S4c**). Mousing over the year 2005 demonstrates that 91% (n=10/11) were 4.3.1 genotypes (pink coloured bars), which declined to 20% (n=8/40) in 2019. Over the same time period, genotype 3.3.2 (mid-blue coloured bars) persisted at low prevalence, and genotypes 2.0.1 and 2.3.3 (green bars) emerged and proliferated. Viewing the ‘Resistance frequencies within genotypes’ plot with default settings (data view as ‘Percentage within genotype’) demonstrates that MDR was only present in genotype 4.3.1.1 (>73%), whereas CipNS was prevalent among all genotypes displayed (**Fig S4d**). Selecting the local Bangladesh genotype ‘4.3.1.3.Bdq’ (**Fig S4d**) from the dropdown menu, one can see this variant has a distinct AMR profile with resistance to ciprofloxacin, ampicillin, sulfonamides and tetracycline, but susceptibility to the older drugs chloramphenicol and trimethoprim as well as ceftriaxone. The ‘Resistance determinants within genotypes’ plot, when viewed with default settings (‘Select drug class’ set to ‘Ciprofloxacin’ and ‘Data view’ set to ‘Percentage per genotype’) reveals that most genotypes harbour a single QRDR mutation (yellow, resulting in CipNS), except for genotype 4.3.1.3.Bdq genomes which have acquired both a QRDR mutation and a *qnrB* gene (purple, resulting in CipR; **Fig S4e**). Despite being CipR, 4.3.1.3.Bdq strains do not appear to have replaced other co-circulating CipNS strains, with both 4.3.1.3.Bdq and CipR remaining at frequencies of <18% between 2005- 2019 (**Fig S4c**). It is tempting to speculate that this may be related in some way to the variant’s susceptibility to once commonly prescribed drugs chloramphenicol and trimethoprim, although it could also be due to a fitness cost associated with the IncFIB(K) plasmid it carries [58,59].

**Figure 4:**
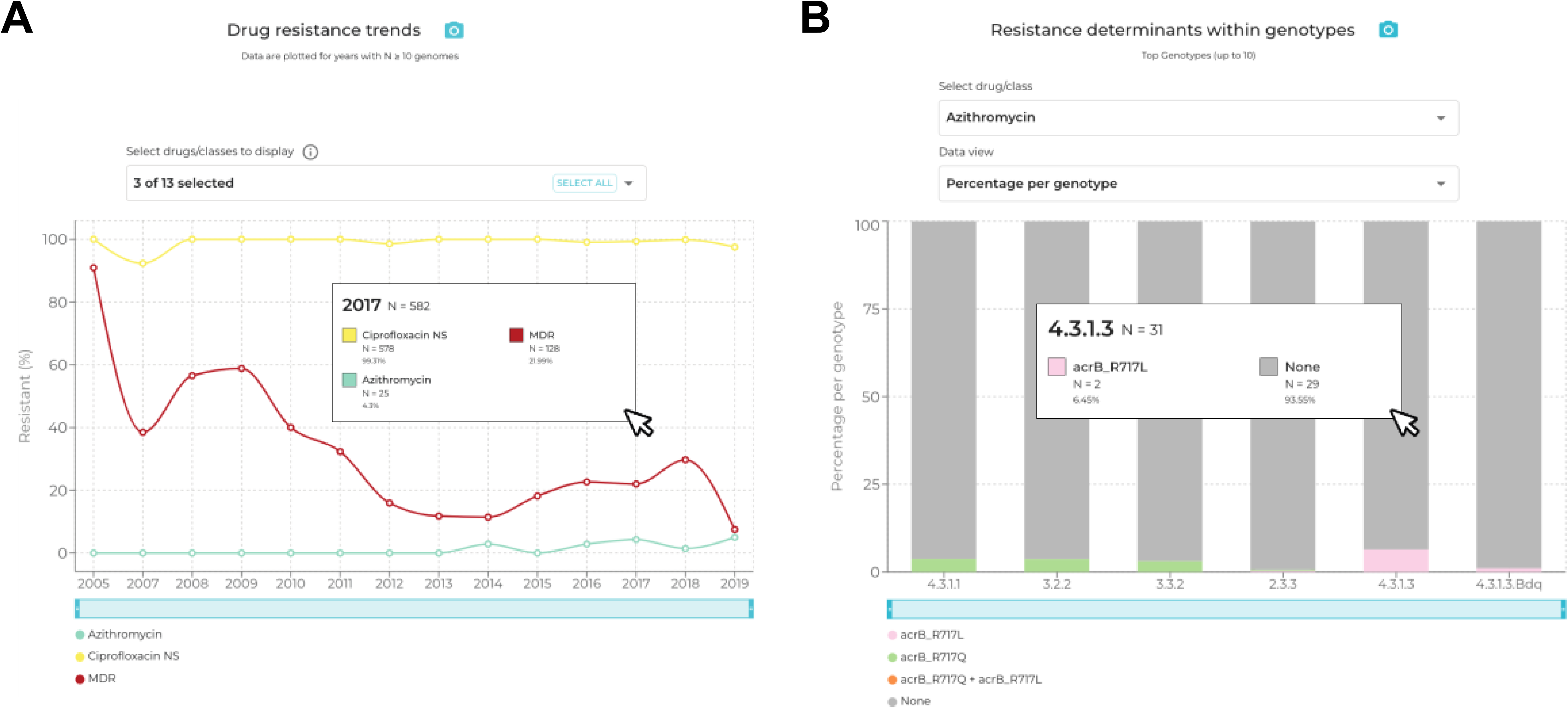
Azithromycin resistance emergence in Bangladesh is associated with different mutations in AcrB, arising in at least six different genotype backgrounds. **(A)** Resistance trends plot showing that ciprofloxacin non-susceptibility has remained near-universal since 2005, while MDR declined. Azithromycin emerged circa 2014, reaching 4-5% in 2017-2019. **(B)** Resistance determinants plot shows that two different types of azithromycin resistance mutations were detected (AcrB-R717L and AcrB-R717Q), in six different genotypes.

The ‘Drug resistance trends’ plot also demonstrates the emergence of azithromycin resistance from 2014 onwards (**Fig 4a**, **Fig 4b**; ≤5% of isolates per year up until 2019) in Bangladesh. The ‘Resistance frequencies within genotypes plot’, with default settings, shows that azithromycin resistance does not appear to be associated with any specific genotype and is occurring in multiple genotype backgrounds (**Fig 4b, Fig S4f**). The molecular mechanisms driving azithromycin resistance among different pathogen genotypes can be viewed in the ‘Resistance determinants within genotypes’ plot by using the ‘Select drug class’ dropdown menu to select ‘Azithromycin’. This reveals the mechanism is a mix of non-synonymous mutations at *acrB* codon 717, with *acrB*-R717Q found in four genotypes and *acrB*-R717L in three genotypes (**Fig 4b, Fig S4f**), including a single example of each mutation in genotype 2.3.3. Examining the global distribution of azithromycin resistance on the map, by selecting ‘Azithromycin resistant’ from the ‘map view’ dropdown menu, highlights that the burden of resistant strains, at present, is low and largely concentrated among South Asian countries (**Fig S5**).

TyphiNET captures several key aspects of the population dynamics and evolution of AMR among Typhi populations in Bangladesh observed across multiple surveillance studies [40,53,54,58,59,70]. These include (i) the decline of MDR over the last two decades, coinciding with a decline in 4.3.1 genotypes; (ii) a sustained high frequency of CipNS cases driven by a diverse range of pathogen genotypes, mostly carrying a single QRDR mutation; (iii) the continued presence of CipR genotype 4.3.1.3.Bdq; and (iv) the timeframe and molecular mechanisms driving the emergence of azithromycin non-susceptibility due to mutations in *acrB* across multiple pathogen genotypes [40,53,54,70]. Fortunately, there does not yet appear to be local establishment or geographical spread of any specific azithromycin-resistant variants (**Fig S5**), however the aggregation of genomic data from multiple sources in the TyphiNET dashboard could facilitate identifying and tracking the emergence of such clones in future.

### Case study 3: Replacement of susceptible genotypes in Malawi with MDR genotype 4.3.1.1

Choosing ‘Multidrug resistant’ from the map dropdown menu shows a few countries with MDR prevalence exceeding 50% (dark red), including Malawi. Setting the time window to 2010 onwards and hovering the mouse cursor over Malawi shows the estimated MDR prevalence for this period is 93% (**Fig S6a**). With the country set to Malawi, the travel filter set to ‘All’ (i.e., including local and travel data) and year range from 2010 to 2019, the ‘Drug resistance trends’ plot shows that MDR prevalence (deep red line; **Fig S6b**) has risen steeply over this decade (this line can be seen more clearly by de-selecting ‘Trimethoprim-sulfamethoxazole’ from the ‘Drugs view’ menu). The proportion amongst the 19 isolates sequenced in 2010 was relatively low (21%), but reached 96% (n=25/26) in 2012 and has been persistently high (>95%) since then. The ‘Genotype distribution’ plot shows that over the same period there was clonal replacement, with the diverse genotypes that were present in 2010 being replaced by genotype 4.3.1.1.EA1 (light pink bars; **Fig S6c**). Hovering the mouse over the graph shows that 4.3.1.1.EA1 prevalence rose from 21% in 2010 to 96% in 2012, and has remained above 95% ever since (**Fig S6c**). The ‘Resistance frequencies within genotypes’ plot demonstrates that resistance to first line drugs is almost entirely associated with genotype 4.3.1.1.EA1 (**Fig S6d**).

The ‘Drug resistance trends’ plot from 2010-2019 (**Fig S6b**) demonstrated that CipNS was relatively low throughout this time period (<16%; yellow line). However, there are two distinct periods within this time frame in which CipNS strains are observed; 2010-2012 and 2018-2019. By adjusting the start and end years to 2010 and 2012, respectively, using the dropdown menus in the top panel, it is apparent from the ‘Resistance frequencies within genotypes’ plot that CipNS during this early period was due to sporadic infections with CipNS genotypes 4.3.1.1 (n=4) and 4.3.1.2 (n=2) (**Fig 5a, S6e**). The ‘Resistance determinants within genotypes’ plot (with ‘Select drug class’ set to ‘Ciprofloxacin’, and ‘Data view’ set to ‘Number of genomes’) shows that genomes from both these genotypes carry one QRDR mutation (**Fig S6f**). However, when viewing the later period of CipNS strains (by adjusting start and end years to 2018 and 2019, respectively) the same two plots reveal that more recent CipNS cases are driven by the emergence of 1–2 QRDR mutations in the locally dominant genotype 4.3.1.1.EA1 (**Figs 5b, S6g-h**). Strains that have acquired two QRDR mutations are of particular concern in this setting due to further elevating ciprofloxacin MIC, which is associated with increases in both fever clearance times and risk of clinical failures [28,49,59,71].

**Figure 5:**
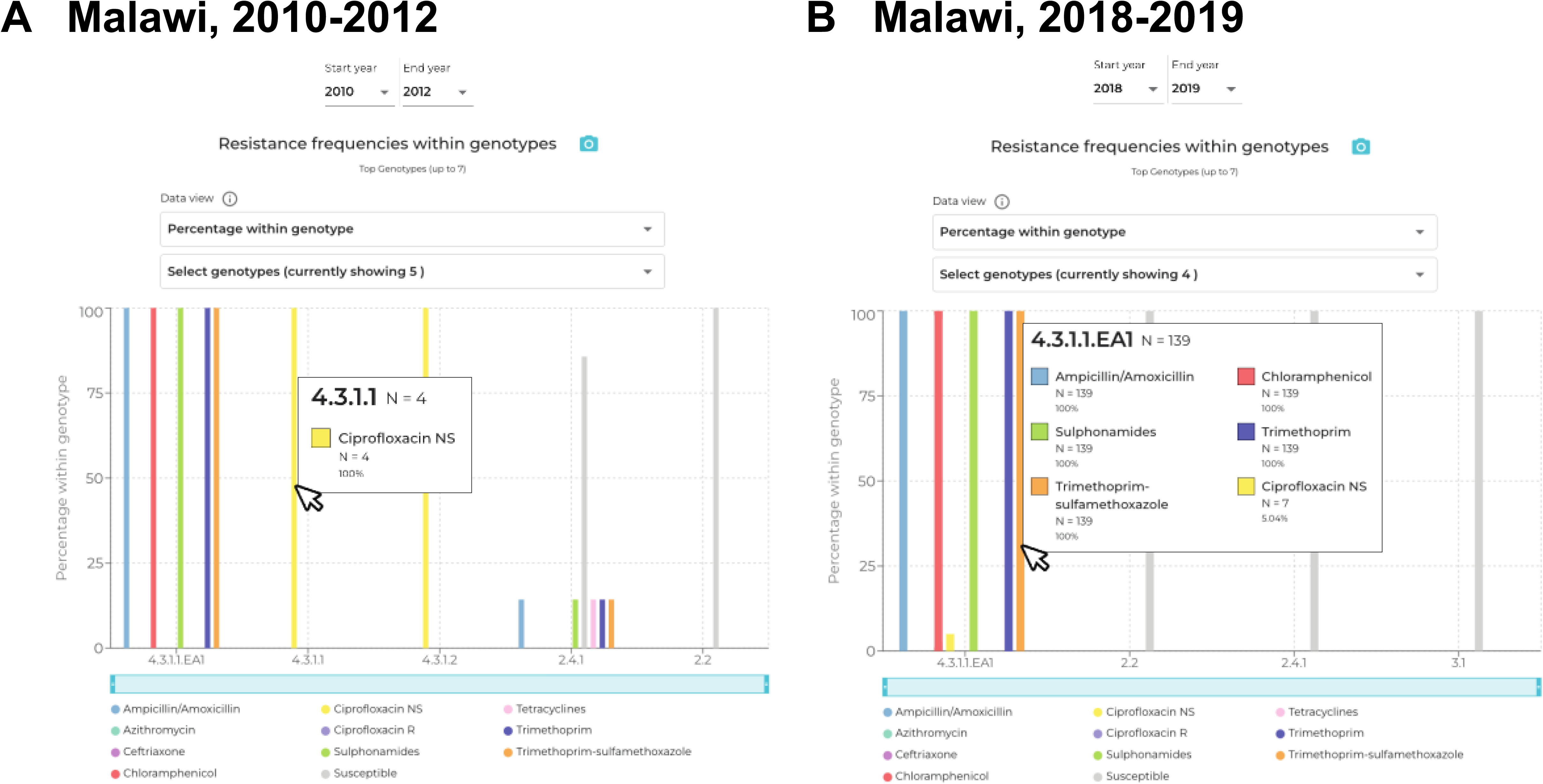
Genotypes associated with Ciprofloxacin non-susceptibility in Malawi in different periods. **(A)** In 2010-2012, CipNS was detected in n=4 isolates of genotype 4.3.1.1 and n=2 isolates of 4.3.1.2, which were susceptible to other drugs. **(B)** In 2018-2019, CipNS emerged in the MDR genotype 4.3.1.1.EA1 (n=7 isolates).

The TyphiNET dashboard highlights the problem of MDR typhoid in Malawi, consistent with several recent genetic and phylogenetic studies [53,72,73]. These include (i) the time of MDR typhoid emergence; (ii) that MDR typhoid is associated with clonal replacement of local genotypes by 4.3.1.1.EA1, which now causes the majority of infections in this setting [53,72]; and (iii) that CipNS is emerging in this setting, driven by the evolution of QRDR mutations in the now-endemic 4.3.1.EA1 strains [73]. Malawi has recently launched a national TCV immunisation program, motivated in large part by the sustained resistance to first-line drugs, and it will be important to monitor the impact of this on the pathogen population. Given the clinical relevance of ciprofloxacin, it will be particularly important to monitor resistance in Malawi and neighbouring countries, which will be facilitated by local and international sequencing efforts and data aggregation in TyphiNET.

### Limitations and future directions

The case studies presented here highlight how TyphiNET can be used to explore the Typhi populations in certain countries, capturing key aspects about the emergence and spread of AMR variants that are supported by current literature. However, the platform is necessarily limited by the availability of source data. India, Pakistan, Bangladesh, Nepal, and Malawi are among the countries with the most Typhi genome data available (n=2327, n=1526, n=1664, n=1300, n=568, respectively), however, this is mainly from a handful of research studies [53,54,58,61,72,73] that may not be representative of typhoid fever in each country. Indeed one use of the dashboard is to highlight data gaps, which may help to prioritise areas for new typhoid surveillance. Routine sequencing of travel-associated cases in the UK and US contributed appreciable data for Pakistan (n=32-217 per year) and Bangladesh (n=15-47 per year), but none from Malawi and very few from Africa in general [22,23]. The importance of pathogen sequencing in supporting infectious disease surveillance and public health response is increasingly recognised, including launch of the WHO-supported International Pathogen Surveillance Network (IPSN) [74] and the 2022-2032 WHO global genomic surveillance strategy for pathogens with pandemic and epidemic potential [75]. Regional and national efforts to strengthen WGS capacity in typhoid-endemic countries include Africa CDC’s Pathogen Genomics Initiative, regional PulseNet networks, and inclusion of WGS in national AMR surveillance in countries such as Pakistan, India, and the Philippines [76–78]. It can therefore be anticipated that the rate of Typhi sequencing is likely to increase, and the representation of cases from endemic areas will improve over time. However, the future utility of TyphiNET and other data aggregation efforts will depend on (i) the WGS data being shared, in a timeframe that is useful to inform and guide decision-making; (ii) the WGS data being accompanied by sufficient and accurate contextual metadata to be useful for epidemiological purposes; (iii) continuous curation of the AMR database hosted at Pathogenwatch; and (iv) ongoing development in response to stakeholder and community feedback. To support these needs, the GTGC was formed in 2021 to support and standardise generation, analysis and sharing of Typhi genome data. A contextual metadata template was developed that includes fields to capture the purpose of sampling as well as country of origin for travel-associated cases [49], and this has been used by the Consortium to curate public genome data for Pathogenwatch and TyphiNET. However, the ongoing relevance of public genome data, and downstream tools that utilise it such as TyphiNET, will depend on metadata standards and data sharing practices being adopted widely by those generating pathogen genomic surveillance data. We hope that this first version of TyphiNET illustrates a potential benefit of such data sharing, that might help encourage such practices. In the meantime, planned future developments to the dashboard include the addition of uncertainty measures (in addition to current visuals, which facilitate understanding limitations of the data by showing the sample size and count data for all data points plotted), and planned activities of the GTGC include aggregation, curation and linkage of AST data to support the ongoing accuracy of AMR predictions from phenotype, as new mechanisms and determinants emerge. Stakeholder engagement and feature development of TyphiNET, as well as extension to other typhoidal and non-typhoidal pathogens, is also ongoing with funding support under the AMRnet project [79].

## Conclusions

Against a backdrop of increasing AMR and vaccine rollouts in multiple countries, the TyphiNET dashboard provides an online resource for monitoring global and national genome-derived trends in AMR and pathogen variants without bioinformatics expertise. These data are potentially informative for implementing and monitoring vaccination and empirical treatment policies for typhoid fever, as well as understanding local variant transmission, enabling improved targeting of costly WASH interventions. As more Typhi WGS data become publicly available and curated by the GTGC, the TyphiNET database will be updated to provide a contemporary overview of ongoing and emerging trends in the pathogen population. Finally, the GTGC and TyphiNET approach provides a useful model for making pathogen genome-derived data broadly accessible that could be applied to other priority pathogens.

## Availability and requirements

**Project name:** TyphiNET AMR surveillance dashboard

**Project home page:** https://www.typhi.net

**Code repository:** https://github.com/typhoidgenomics/TyphiNET/

**Operating system(s):** Platform independent

**Programming language(s):** JavaScript, CSS, HTML

**Other requirements:** None

**License:** GNU GPL-3.0

**Any restrictions to use by non-academics:** None

## List of abbreviations

AMR: Antimicrobial Resistance
API: Application Programming Interface
AST: Antimicrobial Susceptibility testing
AziR: Azithromycin Resistant
BIGSdb: Bacterial Isolate Genome Sequence Database
CefR: Ceftriaxone resistant
cgMLST: core genome multilocus sequence typing
CipNS: Ciprofloxacin Non-susceptible
CipR: Ciprofloxacin Resistant
DDBJ: DNA Data Bank of Japan
EMBL-EBI: European Molecular Biology Laboratory - European Bioinformatics Institute
ENA: European Nucleotide Archive
GLASS: Global Antimicrobial Resistance and Use Surveillance System
GNU-GPL: General User Licence/General Public Licence
GTGC: Global Typhoid Genomics Consortium
H58: Haplotype 58 (genotype 4.3.1)
INSDC: International Nucleotide Sequence Database Collaboration
LMIC: Low- to Middle-Income Country
MDR: Multidrug Resistant
MERN: MongoDB, Express, React, Node
MIC: Minimal Inhibitory Concentration
NCBI: National Center for Biotechnology Information
NITAG: National Immunisation Technical Advisory Groups
PNG: Portable Network Graphics
PDF: Portable Documents Format
QRDR: Quinolone Resistance Determining Region
SEAP: Surveillance for Enteric Fever in Asia Project
STRATAA: Strategic Typhoid Alliance across Africa and Asia
SNP: Single Nucleotide Polymorphism
TCV: Typhoid conjugate vaccine
XDR: Extensively Drug Resistant
WGS: Whole Genome Sequencing
WHO: World Health Organization

## Declarations

### Ethics approval and consent to participate

Not applicable as the project concerns only data which are already in the public domain (please see **Availability of data and materials**), and no personal or clinical information relating to the bacterial isolates is included.

### Consent for publication

Not applicable.

### Availability of data and materials

The TyphiNET dashboard concerns aggregating, processing and visualising genome sequencing data that is already in the public domain. Individual sample accession numbers for the sequences in the European Nucleotide Archive are given in **Supplementary table 1** and are also available in the database download (CSV file) that can be obtained from the dashboard site (https://www.typhi.net).

## Authors’ contributions

KEH and ZAD designed and supervised the study. ZAD, KEH, and MEC analysed the data. LC developed the software. VS tested and improved the map view software features. ZAD wrote the first draft of the manuscript. All authors, including Global Typhoid Genomics Consortium members, reviewed and edited the manuscript. All authors read and approved the final manuscript.

## Acknowledgements

Marie Anne Chattaway is affiliated to the National Institute for Health Research Health Protection Research Unit (NIHR HPRU) in Genomics and Enabling Data at University of Warwick in partnership with the UK Health Security Agency (UKHSA), in collaboration with University of Cambridge and Oxford. Marie Anne Chattaway is based at UKHSA. The views expressed are those of the author(s) and not necessarily those of the NIHR, the Department of Health and Social Care or the UK Health Security Agency.

## Disclaimer

The findings and conclusions of this report are those of the authors and do not necessarily represent the official position of the Centers for Disease Control and Prevention (CDC)

## Global Typhoid Genomics Consortium Group Authorship

David M. Aanensen^1^, Ali H. Abbas^2^, Antoine Abou Fayad^3^, Ayorinde O. Afolayan^4^, Niyaz Ahmed^5^, Irshad Ahmed^6^, Afreenish Amir^7^, Sadia Andleeb^8^, Silvia Argimón^1^, Abraham Aseffa^9^, Philip M. Ashton^10^, Mabel K. Aworh^11,12^, Ashish R. Bavdekar^13^, Marie A. Chattaway^14^, Ka Lip Chew^15^, John A. Crump^16^, Thomas C. Darton^17^, Paula L. Diaz^18^, Christiane Dolecek^19^, Nicholas A. Feasey^20,21^, Andrew R. Greenhill^22^, Madhu Gupta^23^, Mochammad Hatta^24^, Rene S. Hendriksen^25^, Robert S. Heyderman^26^, Odion O. Ikhimiukor^4^, Aamer Ikram^27^, Danielle J. Ingle^28^, Arti Kapil^29^, Jacqueline A. Keane^30^, Karen H. Keddy^31^,Robert A. Kingsley^32^, Myron M. Levine^33^, Calman A. MacLennan^34^, Mailis Maes^30,35^, Jaspreet Mahindroo^36^, Tapfumanei Mashe^37^, Masatomo Morita^38^, Elli Mylona^30^, Geetha Nagaraj^39^, Satheesh Nair^14^, Take K. Naseri^40^, Elisabeth Njamkepo^41^, Sophie Octavia^42^, Iruka N. Okeke^4^, Michael Owusu^43^, Maria Pardos de la Gandara^41^, Andrew J. Pollard^44^, Sadia I. A. Rahman^45^, Saikt Rahman^46^, David A. Rasko^47^, Elrashdy M. Redwan^48,49,50^, Assaf Rokney^51^, Priscilla Rupali^52^, Jean Pierre Rutanga^53^, Jivan Shakya^54^, Senjuti Saha^55^, Michael J. Sikorski^33, 47^, Anthony M. Smith^56, 57^, Kaitlin A. Tagg^58^, Neelam Taneja^59^, Dipesh Tamrakar^60^, Paul Turner^61,19^, James E. Ussher^62^, Sandra Van Puyvelde^30,63^, Koen Vandelannoote^64^, François-Xavier Weill^41^, Vanessa K. Wong^65^, and Jackie Wright^66^

## Affiliations

1. Centre for Genomic Pathogen Surveillance, Pandemic Sciences Institute, University of Oxford, old Road Campus, Oxford, UK

2. Department of Microbiology, Faculty of Veterinary Medicin, University of Kufa, Najaf, 54003, Iraq

3. Department of Experimental Pathology, Immunology and Microbiology, Faculty of Medicine, American University of Beirut, Bliss Street, Beirut, Lebanon

4. Department of Pharmaceutical Microbiology, Faculty of Pharmacy, University of Ibadan, Ibadan, 200284, Oyo State, Nigeria

5. University of Hyderabad, Department of Biotechnology and Bioinformatics, Hyderabad 500046 Telangana State, India

6. Food Safety Laboratory section, National Health Laboratories, Doha, Qatar.

7. National Institutes of Health, Islamabad, Pakistan

8. Department of Industrial Biotechnology, Atta-ur-Rahman School of Applied Biosciences, National University of Sciences and Technology (NUST), Islamabad, Pakistan.

9. Armauer Hansen Research Institute (AHRI), Jimma Road, ALERT Campus, Addis Ababa, POB 1005, Ethiopia

10. Institute of Infection, Veterinary and Ecological Sciences, University of Liverpool, Liverpool L69 3BX, UK

11. Department of Population Health & Pathobiology, College of Veterinary Medicine, North Carolina State University, Raleigh, NC 27607, North Carolina, USA

12. Nigeria Field Epidemiology & Laboratory Training Program, 50 Hailie Selassie Street, Asokoro, Abuja, Nigeria

13. KEM Hospital Research Centre, Pune, 411011, India

14. United Kingdom Health Security Agency, Gastrointestinal Bacteria Reference Unit, 61 Colindale Avenue, London, NW9 5HT, UK

15. National University Hospital, 5 Lower. Kent Ridge Road, Singapore, 119074, Republic of Singapore

16. Centre for International Health, University of Otago, 55 Hanover Street, Dunedin Central, Dunedin 9054, New Zealand

17. Clinical Infection Research Group, Division of Clinical Medicine, School of Medicine & Population Health, University of Sheffield Medical School, Beech Hill Road, Sheffield, S10 2RX, UK

18. Grupo de Microbiología,Instituto Nacional de Salud,Colombia; Avenida Calle 26 · 51- 20, Bogotá ;111321; Colombia

19. Centre for Tropical Medicine and Global Health, University of Oxford, Oxford, United Kingdom

20. School of Medicine, University of St Andrews, St Andrews, Scotland

21. Malawi Liverpool Wellcome Programme, Kamuzu University of Health Sciences, Blantyre, Malawi

22. Federation University Australia, Northways Rd, Churchill, 3842, Victoria, Australia

23. Postgraduate Institute of Medical Education and Research, sector 12, Chandigarh, 160012, India

24. Hasanuddin University, Jl. Perintis Kemerdekaan, Km 10. Makassar, 90245, Indonesia

25. Technical University of Denmark, National Food Institute, Research Group of Global Capacity Building, Kemitorvet, 2800 Kgs. Lyngby, Denmark

26. Division of Infection and Immunity, University College London, Cruciform Building, Gower Street, London, WC1E 6BT, United Kingdom

27. Pakistan Academy of Sciences, Constitution Avenue, Islamabad, 44000, Pakistan

28. Department of Microbiology and Immunology, The University of Melbourne at The Peter Doherty Institute for Infection and Immunity, Melbourne, 3010 Australia.

29. Dept. of Microbiology, AIIMS,New Delhi-110029, India

30. Cambridge Institute of Therapeutic Immunology and Infectious Disease, Department of Medicine, University of Cambridge, Cambridge Biomedical Campus, Cambridge, CB2 0AW, UK

31. Department of Veterinary Tropical Diseases, Faculty of Veterinary Science, University of Pretoria, Old Soutpan Road, Onderstepoort, 0110, South Africa

32. Quadram Institute Bioscience, Rosalind Franklin Road, Norwich Research Park, Norwich, NR47UQ

33. Center for Vaccine Development and Global Health, University of Maryland School of Medicine, 685 W. Baltimore St., Baltimore, MD 21201, USA

34. Jenner Institute, Nuffield Department of Medicine, University of Oxford, Old Road Campus Research Building, Roosevelt Drive, Headington, Oxford, OX3 7DQ, UK

35. Wellcome Sanger Institute, Wellcome Genome Campus, Hinxton, UK

36. MRC Centre for Global Infectious Disease Analysis, School of Public Health, Imperial College London, London, W12 0BZ, UK

37. One Health Office, Ministry of Health and Child Care, Harare, Zimbabwe and Health System Strengthening Unit, World Health Organization, Harare, Zimbabwe

38. Department of Bacteriology I, National Institute of Infectious Diseases, Toyama 1-23- 1, Shinjuku-ku, Tokyo 162-8640, Japan

39. Central Research Laboratory, KIMS, Bengaluru, 560070, INDIA

40. General Practitioner Family Health Clinic, Public Health Specialist & Lead Consultant Naseri & Associates Consultancy, Apia, Samoa

41. Institut Pasteur, Université Paris Cité, 28 rue du Dr Roux, Paris, F-75015, France

42. School of Biotechnology and Biomolecular Sciences, University of New South Wales, Randwick NSW 2052, Australia

43. Kwame Nkrumah University of Science and Technology, Asuogya road, Kumasi, Ghana

44. Oxford Vaccine Group, Department of Paediatrics, University of Oxford, and the NIHR Oxford Biomedical Research Centre, Oxford, OX3 9DU, UK

45. International Centre for Diarrhoeal Disease Research, Bangladesh 68, Shaheed Tajuddin Ahmed Sarani, Mohakhali, Dhaka, 1212, Bangladesh

46. Institute for developing Science and Health initiatives (ideSHi), 167/24, Blue Moon Gram Tower (Lift-11), Kalshi Road, ECB Chattar, Dhaka Cantonment, Dhaka, 1206, Bangladesh

47. Institute of Genome Sciences, University of Maryland School of Medicine, 670 W. Baltimore Street, Baltimore, MD 21201, USA

48. King Abdulaziz University, Faculty of Science, Department Biological Science , Jeddah 21589, Saudi Arabia

49. Centre of Excellence in Bionanoscience Research, King Abdulaziz University, Jeddah 21589, Saudi Arabia

50. Therapeutic and Protective Proteins Laboratory, Protein Research Department, Genetic Engineering and Biotechnology Research Institute, City of Scientific Research and Technological Applications (SRTA-City), New Borg EL-Arab, 21934, Alexandria, Egypt

51. Ministry of Health, Ya’akov Eliav 9, Jerusalem, 91342, Israel

52. Christian Medical College Vellore, Ida Scudder Road, Vellore, Tamilnadu, 632004, India

53. College of Science and Technology (CST), University of Rwanda, KN 7 Ave, Kigali, Rwanda

54. Mycobacterial Research Laboratories (MRL), Anandaban Hospital, Lele, Lalitpur, Nepal

55. Child Health Research Foundation, 23/2 Khilji Road, Mohammadpur, Dhaka, 1207, Bangladesh

56. Centre for Enteric Diseases, National Institute for Communicable Diseases, Division of the National Health Laboratory Service, 1 Modderfontein Road, Johannesburg, 2000, South Africa

57. Department of Medical Microbiology, School of Medicine, Faculty of Health Sciences, University of Pretoria, Lynnwood Road, Pretoria, 0002, South Africa

58. Centers for Disease Control and Prevention, 1600 Clifton Rd, Atlanta, GA, 30333, USA,

59. Department of Medical Microbiology, Postgraduate Institute of Medical Education and Research, Chandigarh, Pin 160012, India

60. Dhulikhel Hospital, Kathmandu University Hospital, Dhulikhel, Nepal

61. Cambodia-Oxford Medical Research Unit, Angkor Hospital for Children, Siem Reap, Cambodia

62. Department of Microbiology and Immunology, University of Otago, 720 Cumberland St, Dunedin, 9054, New Zealand

63. University of Antwerp, Universiteitsplein 1, 2000 Antwerp, Belgium

64. Institut Pasteur du Cambodge, 5 Blvd Monivong, Phnom Penh, Cambodia

65. Gastrointestinal Infections and Food safety (One Health) Division, UKHSA, 61 Colindale Avenue, NW9 5EQ, London, UK

66. Institute of Environmental Science and Research, 27 Creyke Road, Ilam, Christchurch 8041, New Zealand

## References

1. Meiring JE, Khanam F, Basnyat B, Charles RC, Crump JA, Debellut F, et al. Typhoid fever. Nat Rev Dis Prim. 2023;9:71.

2. GBD 2019 Diseases and Injuries Collaborators, Vos T, Lim SS, Abbafati C, Abbas KM, Abbasi M, et al. Global burden of 369 diseases and injuries in 204 countries and territories, 1990–2019: a systematic analysis for the Global Burden of Disease Study 2019. Lancet. 2020;396:1204–22.

3. GBD 2017 Typhoid and Paratyphoid Collaborators. The global burden of typhoid and paratyphoid fevers: a systematic analysis for the Global Burden of Disease Study 2017. Lancet Infect Dis. 2019;19:369–81.

4. Murti BR, Rajyalakshmi K, Bhaskaran CS. Resistance of *Salmonella* Typhi to chloramphenicol. J Clin Pathol. 1962;15:544.

5. Dyson ZA, Klemm EJ, Palmer S, Dougan G. Antibiotic Resistance and Typhoid. Clin Infect Dis. 2019;68:S165–70.

6. Parry CM, Vinh H, Chinh NT, Wain J, Campbell JI, Hien TT, et al. The Influence of Reduced Susceptibility to Fluoroquinolones in *Salmonella enterica* Serovar Typhi on the Clinical Response to Ofloxacin Therapy. Plos Negl Trop Dis. 2011;5:e1163.

7. Espinoza LMC, McCreedy E, Holm M, Im J, Mogeni OD, Parajulee P, et al. Occurrence of Typhoid Fever Complications and Their Relation to Duration of Illness Preceding Hospitalization: A Systematic Literature Review and Meta-analysis. Clin Infect Dis. 2019;69:S435–48.

8. Butler T, Knight J, Nath SK, Speelman P, Roy SK, Azad MA. Typhoid fever complicated by intestinal perforation: a persisting fatal disease requiring surgical management. Rev Infect Dis. 1985;7:244–56.

9. Crump JA. Progress in Typhoid Fever Epidemiology. Clin Infect Dis. 2019;68:S4–9.

10. Wain J, Diep TS, Bay PVB, Walsh AL, Vinh H, Duong NM, et al. Specimens and culture media for the laboratory diagnosis of typhoid fever. J Infect Dev Ctries. 2008;2:469–74.

11. Mogasale V, Ramani E, Mogasale VV, Park J. What proportion of *Salmonella* Typhi cases are detected by blood culture? A systematic literature review. Ann Clin Microbiol Antimicrob. 2016;15:32.

12. World Health Organization. The WHO AWaRe (Access, Watch, Reserve) antibiotic book. Geneva; 2022.

13. Andrews JR, Baker S, Marks F, Alsan M, Garrett D, Gellin BG, et al. Typhoid conjugate vaccines: a new tool in the fight against antimicrobial resistance. Lancet Infect Dis. 2019;19:e26–30.

14. World Health Organization. Typhoid vaccines: WHO position paper. Week Epidemiol Rec. 2018;13:153–72.

15. Yousafzai MT, Karim S, Qureshi S, Kazi M, Memon H, Junejo A, et al. Effectiveness of typhoid conjugate vaccine against culture-confirmed *Salmonella enterica* serotype Typhi in an extensively drug-resistant outbreak setting of Hyderabad, Pakistan: a cohort study. Lancet Glob Health. 2021;9:e1154–62.

16. Lightowler MS, Manangazira P, Nackers F, Herp MV, Phiri I, Kuwenyi K, et al. Effectiveness of typhoid conjugate vaccine in Zimbabwe used in response to an outbreak among children and young adults: A matched case control study. Vaccine. 2022;40:4199– 210.

17. World Health Organization. Pakistan first country to introduce new typhoid vaccine into routine immunization programme [Internet]. 2019 [cited 2023 May 30]. Available from: https://www.emro.who.int/pak/pakistan-news/pakistan-first-country-to-introduce-new-typhoid-vaccine-into-routine-immunization-programme.html

18. Oberman E. Typhoid conjugate vaccine arrives in Zimbabwe [Internet]. 2021 [cited 2023 May 30]. Available from: https://www.coalitionagainsttyphoid.org/typhoid-conjugate-vaccine-arrives-in-zimbabwe/

19. Mashe T, Leekitcharoenphon P, Mtapuri-Zinyowera S, Kingsley RA, Robertson V, Tarupiwa A, et al. *Salmonella enterica* serovar Typhi H58 clone has been endemic in Zimbabwe from 2012 to 2019. J Antimicrob Chemother. 2021;76(5):1375.

20. Thilliez G, Mashe T, Chaibva BV, Robertson V, Bawn M, Tarupiwa A, et al. Population structure of *Salmonella enterica* Typhi in Harare, Zimbabwe (2012–19) before typhoid conjugate vaccine roll-out: a genomic epidemiology study. Lancet Microbe. 2023;4:e1005– 14.

21. World Health Organization. Global antimicrobial resistance and use surveillance system (GLASS) report 2022 [Internet]. Geneva; 2022. Available from: https://www.who.int/publications/i/item/9789240062702

22. Ingle DJ, Nair S, Hartman H, Ashton PM, Dyson ZA, Day M, et al. Informal genomic surveillance of regional distribution of *Salmonella* Typhi genotypes and antimicrobial resistance via returning travellers. Akullian A, editor. PLoS Negl Trop Dis. 2019;13:e0007620.

23. Carey ME, Dyson ZA, Ingle DJ, Amir A, Aworh MK, Chattaway MA, et al. Global diversity and antimicrobial resistance of typhoid fever pathogens: Insights from a meta-analysis of 13,000 *Salmonella* Typhi genomes. eLife. 2023;12:e85867.

24. Ashton PM, Nair S, Peters TM, Bale JA, Powell DG, Painset A, et al. Identification of *Salmonella* for public health surveillance using whole genome sequencing. PeerJ. 2016;4:e1752.

25. Chattaway MA, Gentle A, Nair S, Tingley L, Day M, Mohamed I, et al. Phylogenomics and antimicrobial resistance of *Salmonella* Typhi and Paratyphi A, B and C in England, 2016– 2019. Microb Genom. 2021;7:000633.

26. Dyson ZA, Holt KE. Five Years of GenoTyphi: Updates to the Global *Salmonella* Typhi Genotyping Framework. J Infect Dis. 2021;224:S775–80.

27. Wong VK, Baker S, Connor TR, Pickard D, Page AJ, Dave J, et al. An extended genotyping framework for *Salmonella enterica* serovar Typhi, the cause of human typhoid. Nat Commun. 2016;7:12827.

28. Argimon S, Yeats CA, Goater RJ, Abudahab K, Taylor B, Underwood A, et al. A global resource for genomic predictions of antimicrobial resistance and surveillance of *Salmonella* Typhi at pathogenwatch. Nat Commun. 2021;12:2879–12.

29. Nair S, Chattaway M, Langridge GC, Gentle A, Day M, Ainsworth EV, et al. ESBL- producing strains isolated from imported cases of enteric fever in England and Wales reveal multiple chromosomal integrations of blaCTX-M-15 in XDR *Salmonella* Typhi. J Antimicrob Chemother. 2021;76:1459–66.

30. Watkins LKF, Winstead A, Appiah GD, Friedman CR, Medalla F, Hughes MJ, et al. Update on Extensively Drug-Resistant *Salmonella* Serotype Typhi Infections Among Travelers to or from Pakistan and Report of Ceftriaxone-Resistant *Salmonella* Serotype Typhi Infections Among Travelers to Iraq — United States, 2018–2019. Morbidity Mortal Wkly Rep. 2020;69:618–22.

31. Nabarro LE, McCann N, Herdman MT, Dugan C, Ladhani S, Patel D, et al. British infection association guidelines for the diagnosis and management of enteric fever in England. J Infection. 2022;84:469–89.

32. Gauld JS, Olgemoeller F, Heinz E, Nkhata R, Bilima S, Wailan AM, et al. Spatial and genomic data to characterize endemic typhoid transmission. Clin Infect Dis. 2022;74:ciab745.

33. Waddington C, Carey ME, Boinett CJ, Higginson E, Veeraraghavan B, Baker S. Exploiting genomics to mitigate the public health impact of antimicrobial resistance. Genome Med. 2022;14:15.

34. Armstrong GL, MacCannell DR, Taylor J, Carleton HA, Neuhaus EB, Bradbury RS, et al. Pathogen Genomics in Public Health. New Engl J Med. 2019;381:2569–80.

35. National Center for Biotechnology Information. NCBI Pathogen Detection [Internet]. Available from: https://www.ncbi.nlm.nih.gov/pathogens/isolates/#taxgroup_name:%22Salmonella%20enterica%22

36. Zhou Z, Alikhan N-F, Mohamed K, Fan Y, Group the AS, Achtman M, et al. The EnteroBase user’s guide, with case studies on *Salmonella* transmissions, *Yersinia pestis* phylogeny, and *Escherichia* core genomic diversity. Genome Res. 2020;30:138–52.

37. *Salmonella* EnteroBase [Internet]. Available from: https://enterobase.warwick.ac.uk/species/index/senterica

38. Jolley KA, Maiden MCJ. BIGSdb: Scalable analysis of bacterial genome variation at the population level. BMC Bioinformatics. 2010;11:595.

39. Klemm EJ, Shakoor S, Page AJ, Qamar FN, Judge K, Saeed DK, et al. Emergence of an Extensively Drug-Resistant *Salmonella enterica* Serovar Typhi Clone Harboring a Promiscuous Plasmid Encoding Resistance to Fluoroquinolones and Third-Generation Cephalosporins. mBio. 2018;9.

40. Hooda Y, Sajib MSI, Rahman H, Luby SP, Bondy-Denomy J, Santosham M, et al. Molecular mechanism of azithromycin resistance among typhoidal *Salmonella* strains in Bangladesh identified through passive pediatric surveillance. PLoS Negl Trop Dis. 2019;13:e0007868.

41. Cerdeira L, Sousa M, Dyson Z. lcerdeira/Spyder: Spyder. 2021; Available from: 10.5281/zenodo.4740169

42. Cerdeira L, vLSHTM, Dyson ZA, Holt K. typhoidgenomics/TyphiNET: v1.5.1. 2024; Available from: 10.5281/zenodo.10667321

43. Dyson ZA, Holt KE. Five years of GenoTyphi: updates to the global *Salmonella* Typhi genotyping framework. J Infect Dis. 2021;224:jiab414.

44. Matono T, Morita M, Yahara K, Lee K, Izumiya H, Kaku M, et al. Emergence of Resistance Mutations in *Salmonella enterica* Serovar Typhi Against Fluoroquinolones. Open Forum Infect Dis. 2017;4:ofx230.

45. Ingle DJ, Andersson P, Valcanis M, Wilmot M, Easton M, Lane C, et al. Genomic Epidemiology and Antimicrobial Resistance Mechanisms of Imported Typhoid in Australia. Antimicrob Agents Chemother. 2021;65:e01200–21.

46. Centers for Disease Control and Prevention (CDC). National Typhoid and Paratyphoid Fever Surveillance Overview [Internet]. Atlanta, Georgia: US Department of Health and Human Services, CDC; 2011. Available from: https://www.cdc.gov/ncezid/dfwed/PDFs/typhi_surveillance_overview_508c.pdf

47. Chen J, Long JE, Vannice K, Shewchuk T, Kumar S, Steele AD, et al. Taking on Typhoid: Eliminating Typhoid Fever as a Global Health Problem. Open Forum Infect Dis. 2023;10:S74– 81.

48. Day MR, Doumith M, Nascimento VD, Nair S, Ashton PM, Jenkins C, et al. Comparison of phenotypic and WGS-derived antimicrobial resistance profiles of *Salmonella enterica* serovars Typhi and Paratyphi. J Antimicrob Chemoth. 2018;73:365–72.

49. Wain J, Diep TS, Ho VA, Walsh AM, Hoa NTT, Parry CM, et al. Quantitation of Bacteria in Blood of Typhoid Fever Patients and Relationship between Counts and Clinical Features, Transmissibility, and Antibiotic Resistance. J Clin Microbiol. 1998;36:1683–7.

50. European Committee on Antimicrobial Susceptibility Testing. Clinical breakpoints - breakpoints and guidance [Internet]. 2023 [cited 2024 Feb 25]. Available from: https://www.eucast.org/clinical_breakpoints

51. Ochieng C, Chen JC, Osita MP, Katz LS, Griswold T, Omballa V, et al. Molecular characterization of circulating *Salmonella* Typhi strains in an urban informal settlement in Kenya. Plos Neglect Trop D. 2022;16:e0010704.

52. Davies MR, Duchene S, Valcanis M, Jenkins AP, Jenney A, Rosa V, et al. Genomic epidemiology of *Salmonella* Typhi in Central Division, Fiji, 2012 to 2016. Lancet Regional Heal - West Pac. 2022;24:100488.

53. Dyson ZA, Ashton PM, Khanam F, Chunga A, Shakya M, Meiring J, et al. Genomic epidemiology and antimicrobial resistance transmission of *Salmonella* Typhi and Paratyphi A at three urban sites in Africa and Asia. medRxiv. 2023; 2023.03.11.23286741; doi: 10.1101/2023.03.11.23286741

54. Silva KE da, Tanmoy AM, Pragasam AK, Iqbal J, Sajib MSI, Mutreja A, et al. The international and intercontinental spread and expansion of antimicrobial-resistant *Salmonella* Typhi: a genomic epidemiology study. Lancet Microbe. 2022;3:e567–77.

55. Kariuki S, Keddy KH, Antonio M, Okeke IN. Antimicrobial resistance surveillance in Africa: Successes, gaps and a roadmap for the future. Afr J Lab Med. 2018;7:924.

56. Song Y, Roumagnac P, Weill FX, Wain J, Dolecek C, Mazzoni CJ, et al. A multiplex single nucleotide polymorphism typing assay for detecting mutations that result in decreased fluoroquinolone susceptibility in *Salmonella enterica* serovars Typhi and Paratyphi A. Journal of Antimicrobial Chemotherapy. 2010;65:1631–41.

57. Thanh DP, Karkey A, Dongol S, Thi NH, Thompson CN, Rabaa MA, et al. A novel ciprofloxacin-resistant subclade of H58 *Salmonella* Typhi is associated with fluoroquinolone treatment failure. eLife. 2016;5:e14003.

58. Rahman SIA, Dyson ZA, Klemm EJ, Khanam F, Holt KE, Chowdhury EK, et al. Population structure and antimicrobial resistance patterns of *Salmonella* Typhi isolates in urban Dhaka, Bangladesh from 2004 to 2016. PLoS Negl Trop Dis. 2020;14:e0008036.

59. Tanmoy AM, Westeel E, Bruyne KD, Goris J, Rajoharison A, Sajib MSI, et al. *Salmonella enterica* Serovar Typhi in Bangladesh: Exploration of Genomic Diversity and Antimicrobial Resistance. mBio. 2018;9:e02112–18.

60. Argimón S, Nagaraj G, Shamanna V, Darmavaram S, Vasanth AK, Prasanna A, et al. Circulation of third-generation cephalosporin resistant *Salmonella* Typhi in Mumbai, India. Clin Infect Dis. 2021;74:ciab897.

61. Rasheed F, Saeed M, Alikhan N-F, Baker D, Khurshid M, Ainsworth EV, et al. Emergence of Resistance to Fluoroquinolones and Third-Generation Cephalosporins in *Salmonella* Typhi in Lahore, Pakistan. Microorganisms. 2020;8:1336.

62. Gul D, Potter RF, Riaz H, Ashraf ST, Wallace MA, Munir T, et al. Draft Genome Sequence of a *Salmonella enterica* Serovar Typhi Strain Resistant to Fourth-Generation Cephalosporin and Fluoroquinolone Antibiotics. Genome Announc. 2017;5:e00850–17.

63. Public Health England. UK Public Health Resistance Alert: *Salmonella* Typhi resistant to third generation cephalosporins isolated in England from a traveller returning from Pakistan. Health Protection Report. 2017;11.

64. Eshaghi A, Zittermann S, Bharat A, Mulvey MR, Allen VG, Patel SN. Importation of Extensively Drug-Resistant *Salmonella enterica* Serovar Typhi Cases in Ontario, Canada. Antimicrob Agents Chemother. 2020;64.

65. Chirico C, Tomasoni LR, Corbellini S, Francesco MAD, Caruso A, Scaltriti E, et al. The first Italian case of XDR *Salmonella* Typhi in a traveler returning from Pakistan, 2019: An alert for increased surveillance also in European countries? Travel Med Infect Dis. 2020;36:101610.

66. Liu P-Y, Wang K-C, Hong Y-P, Chen B-H, Shi Z-Y, Chiou C-S. The first imported case of extensively drug-resistant *Salmonella enterica* serotype Typhi Infection in Taiwan and the antimicrobial therapy. J Microbiol Immunol Infect. 2020;54:740–4.

67. Al-Rashdi A, Kumar R, Al-Bulushi M, Abri SA, Al-Jardani A. Genomic analysis of the first cases of extensively drug-resistant, travel-related *Salmonella enterica* serovar Typhi in Oman. Ijid Regions. 2021;1:135–41.

68. Wang Y, Lu D, Jin Y, Wang H, Lyu B, Zhang X, et al. Extensively Drug-Resistant (XDR) Salmonella Typhi Outbreak by Waterborne Infection — Beijing Municipality, China, January– February 2022. China Cdc Wkly. 2022;4:254–8.

69. Carey ME, Jain R, Yousuf M, Maes M, Dyson ZA, Thu TNH, et al. Spontaneous Emergence of Azithromycin Resistance in Independent Lineages of *Salmonella* Typhi in Northern India. Clin Infect Dis. 2021;72:e120–7.

70. Sajib MSI, Tanmoy AM, Hooda Y, Rahman H, Andrews JR, Garrett DO, et al. Tracking the Emergence of Azithromycin Resistance in Multiple Genotypes of Typhoidal *Salmonella*. mBio. 2021;12:e03481–20.

71. Day MR, Doumith M, Nascimento VD, Nair S, Ashton PM, Jenkins C, et al. Comparison of phenotypic and WGS-derived antimicrobial resistance profiles of *Salmonella enterica* serovars Typhi and Paratyphi. J Antimicrob Chemoth. 2017;73:365–72.

72. Feasey NA, Gaskell K, Wong V, Msefula C, Selemani G, Kumwenda S, et al. Rapid emergence of multidrug resistant, H58-lineage Salmonella typhi in Blantyre, Malawi. PLoS Negl Trop Dis. 2015;9:e0003748.

73. Ashton PM, Chirambo AC, Meiring JE, Patel PD, Mbewe M, Silungwe N, et al. Evaluating the relationship between ciprofloxacin prescription and non-susceptibility in *Salmonella* Typhi in Blantyre, Malawi: an observational study. Lancet Microbe. 2024;5:e226–34.

74. World Health Organization. International Pathogen Surveillance Network (IPSN) [Internet]. [cited 2024 Feb 25]. Available from: https://www.who.int/initiatives/international-pathogen-surveillance-network

75. World Health Organization. Global genomic surveillance strategy for pathogens with pandemic and epidemic potential, 2022–2032 [Internet]. Geneva: World Health Organization; 2022. Available from: https://www.who.int/publications/i/item/9789240046979

76. Inzaule SC, Tessema SK, Kebede Y, Ouma AEO, Nkengasong JN. Genomic-informed pathogen surveillance in Africa: opportunities and challenges. Lancet Infect Dis. 2021;21:e281–9.

77. Davedow T, Carleton H, Kubota K, Palm D, Schroeder M, Gerner-Smidt P, et al. PulseNet International Survey on the Implementation of Whole Genome Sequencing in Low and Middle-Income Countries for Foodborne Disease Surveillance. Foodborne Pathog Dis. 2022;19:332–40.

78. Stevens EL, Carleton HA, Beal J, Tillman GE, Lindsey RL, Lauer AC, et al. Use of Whole Genome Sequencing by the Federal Interagency Collaboration for Genomics for Food and Feed Safety in the United States. J Food Protect. 2022;85:755–72.

79. Holt KE. AMRnet [Internet]. 2023 [cited 2023 Jul 2]. Available from: https://www.lshtm.ac.uk/research/centres-projects-groups/amrnet

