## Supplementary materials for "The TyphiNET data visualisation dashboard: Unlocking *Salmonella* Typhi genomics data to support public health"

**Data S1: TyphiNET downloadable report (pdf format) of all data available from Pakistan.**

**Table S1: TyphiNET downloadable line list (csv format) of all genome-derived data presented in TyphiNET.**

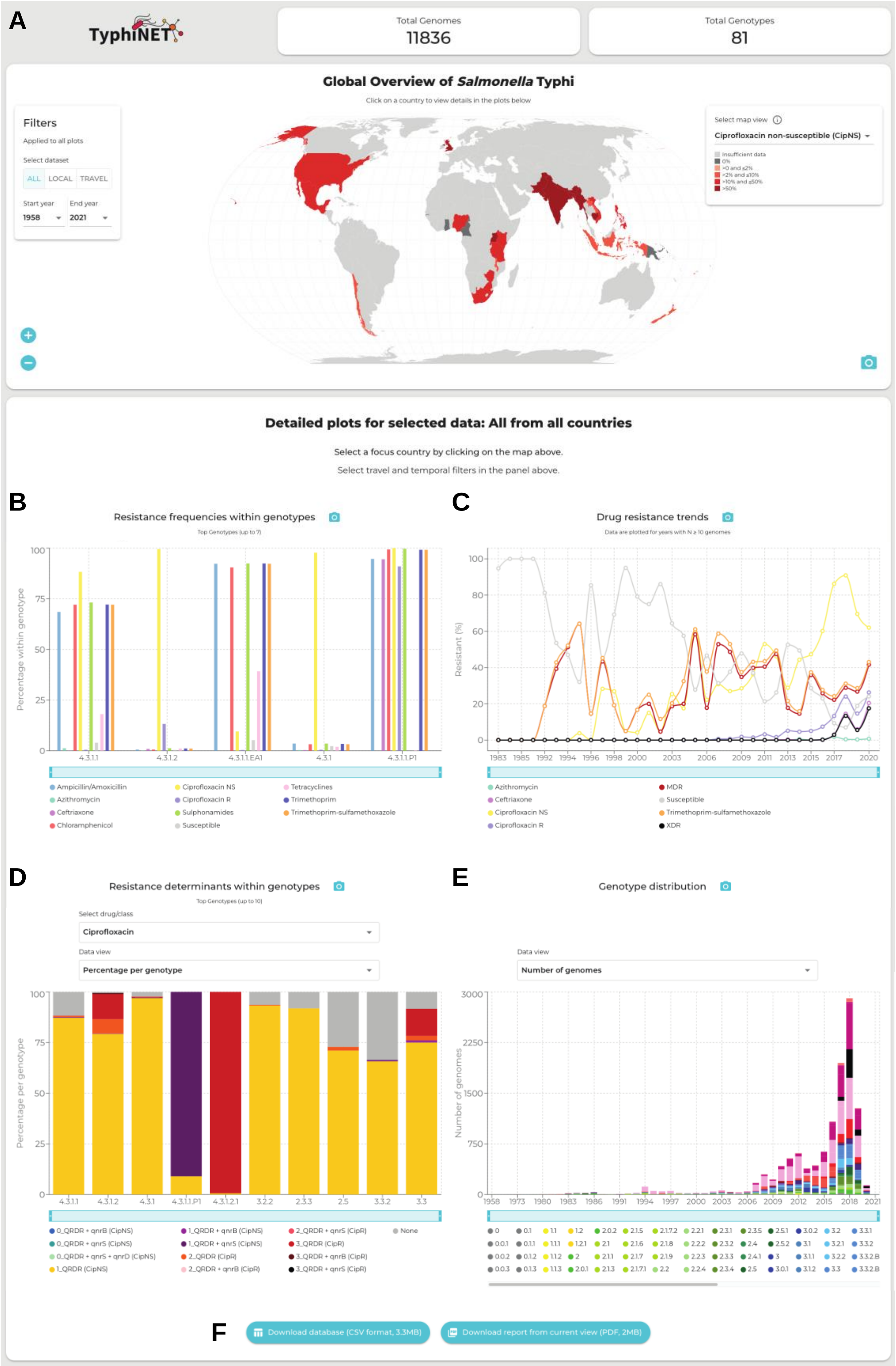
A
TyphiNET
Total Genomes
11836
Total Genotypes
81
Global Overview of Salmonella Typhi
Click on a country to view details in the plots below
Filters
Applied to all plots
Select dataset
ALL
LOCAL
TRAVEL
Start year
1958
End year
2021
Select map view
Ciprofloxacin non-susceptible (CipNS)
Insufficient data
0%
>0 and ≤2%
>2% and ≤10%
>10% and ≤50%
>50%
Detailed plots for selected data: All from all countries
Select a focus country by clicking on the map above.
Select travel and temporal filters in the panel above.
B
Resistance frequencies within genotypes
Top Genotypes (up to 7)
Percentage within genotype
4.3.1.1
4.3.1.2
4.3.1.1.EA1
4.3.1
4.3.1.1.P1
Ampicillin/Amoxicillin
Azithromycin
Ceftriaxone
Chloramphenicol
Ciprofloxacin NS
Ciprofloxacin R
Sulphonamides
Susceptible
Tetracyclines
Trimethoprim
Trimethoprim-sulfamethoxazole
C
Drug resistance trends
Data are plotted for years with N ≥ 10 genomes
Resistant (%)
Azithromycin
Ceftriaxone
Ciprofloxacin NS
Ciprofloxacin R
MDR
Susceptible
Trimethoprim-sulfamethoxazole
XDR
D
Resistance determinants within genotypes
Top Genotypes (up to 10)
Select drug/class
Ciprofloxacin
Data view
Percentage per genotype
Percentage per genotype
4.3.1.1
4.3.1.2
4.3.1
4.3.1.1.P1
4.3.1.2.1
3.2.2
2.3.3
2.5
3.3.2
3.3
0_QRDR + qnrB (CipNS)
0_QRDR + qnrS (CipNS)
0_QRDR + qnrS + qnrD (CipNS)
1_QRDR (CipNS)
1_QRDR + qnrB (CipNS)
1_QRDR + qnrS (CipNS)
2_QRDR (CipR)
2_QRDR + qnrB (CipR)
2_QRDR + qnrS (CipR)
3_QRDR (CipR)
3_QRDR + qnrB (CipR)
3_QRDR + qnrS (CipR)
None
E
Genotype distribution
Data view
Number of genomes
Number of genomes
F
Download database (CSV format, 3.3MB)
Download report from current view (PDF, 2MB)

**Figure S1. Dashboard User Interface. (A) Global map view of ciprofloxacin non-susceptible Typhi.** Countries on the map are coloured by ciprofloxacin non-susceptibility frequency as per the inset legend (top right of map). Data are shown where there are ≥20 sequences available for the country of interest. **(B) Resistance frequencies within genotypes plot shows frequencies of resistance to different drug classes for common genotypes.** Bars are coloured by drug class as per the inset legend. **(C) Drug resistance trends plot shows frequencies of resistance to selected drug classes per year.** Lines are coloured by drug class as per the inset legend. **(D) Resistance determinants within pathogen genotypes plot shows the distribution of specific genes and mutations mediating resistance to a select drug class within common pathogen genotypes.** Bars are coloured by resistance determinants as per the inset legend. **(E) Annual genotype distribution plot shows the frequencies of pathogen genotypes circulating annually.** Bars are coloured by pathogen genotype as per the inset legend. **(F) Downloadable database (csv format) and report (pdf report).**

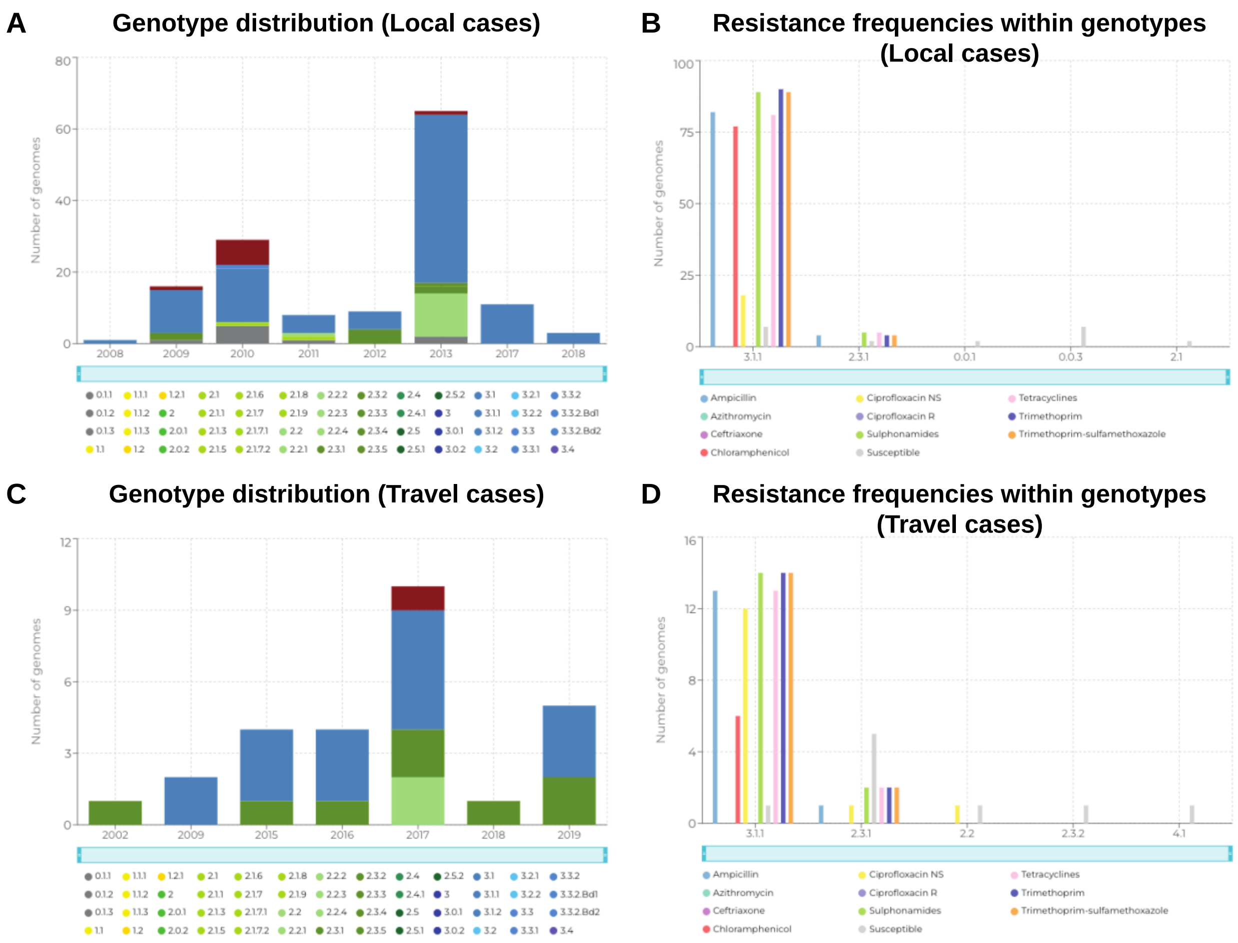


**Figure S2. Frequencies of genotypes and antimicrobial resistance among locally collected and travel-associated cases from Nigeria. (A) National annual pathogen genotype frequencies, derived from locally collected cases.** Bars are coloured by pathogen genotypes as per the inset legend. **(B) Resistance frequencies stratified by pathogen genotype, derived from locally collected cases.** Bars are coloured by drug class as per the inset legend. **(C) National annual pathogen genotype frequencies, derived from travel-associated cases.** Bars are coloured by pathogen genotypes as per the inset legend. **(D) Resistance frequencies stratified by pathogen genotype, derived from travel-associated collected cases.** Bars are coloured by drug class as per the inset legend.

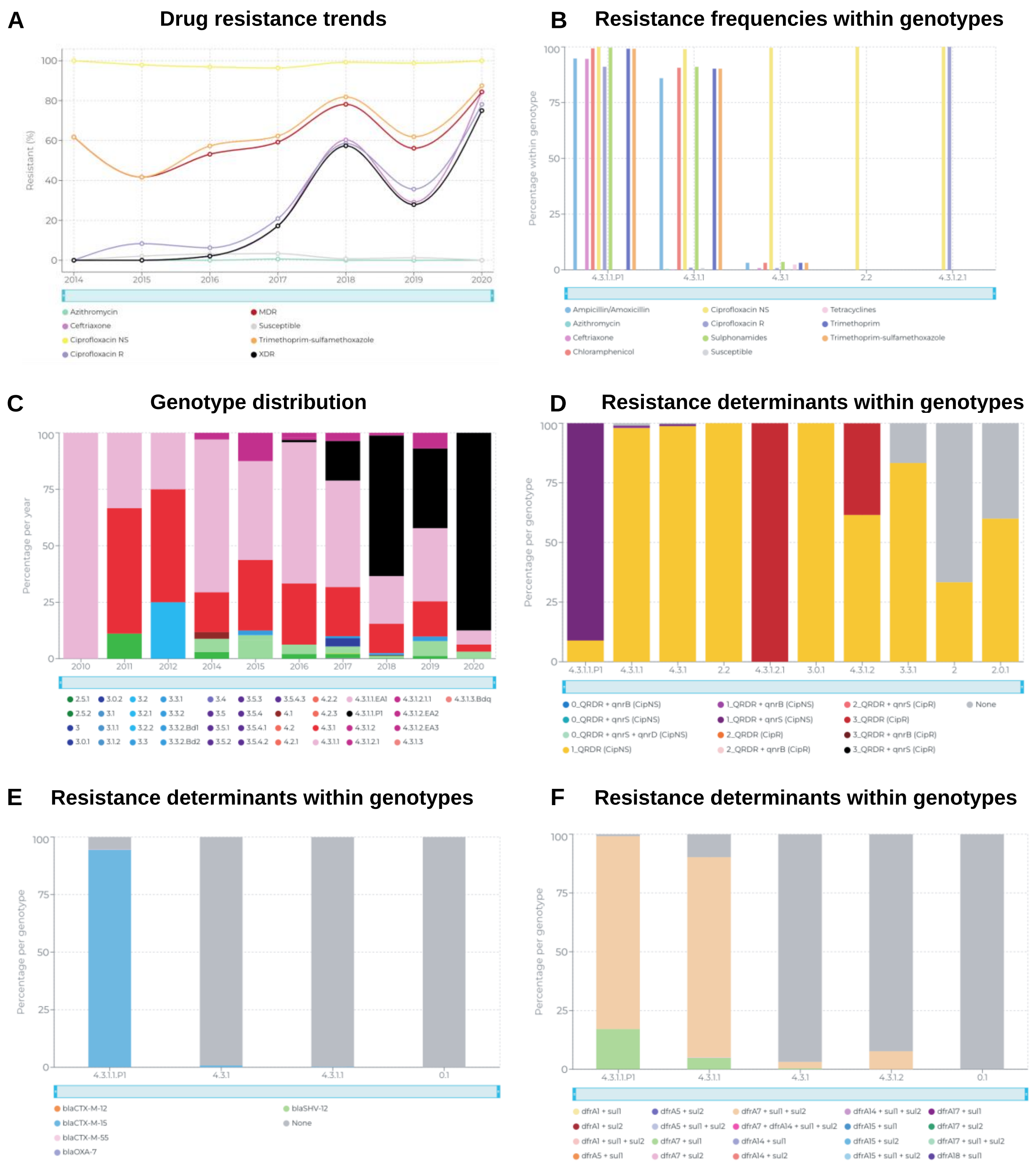


**Figure S3. Exploring the emergence of XDR Typhi in Pakistan with the TyphiNET dashboard. (A) Drug resistance trends over time plot shows frequencies of resistance to selected drug classes in Pakistan per year.** Lines are coloured as per the inset legend. **(B) 'Resistance frequencies within genotypes' plot shows frequencies of resistance to different drug classes, within common genotypes circulating in Pakistan.** Bars are coloured according to the inset legend. **(C) 'Annual genotype distribution' plot shows the frequencies of pathogen genotypes**

**circulating in Pakistan per year.** Genotypes are coloured as per the inset legend. **(D,E,F) 'Resistance determinants within genotypes' plots show the distribution of specific genes and mutations mediating resistance to a selected drug class, within common pathogen genotypes.** Bars are coloured as per the inset legend**.**

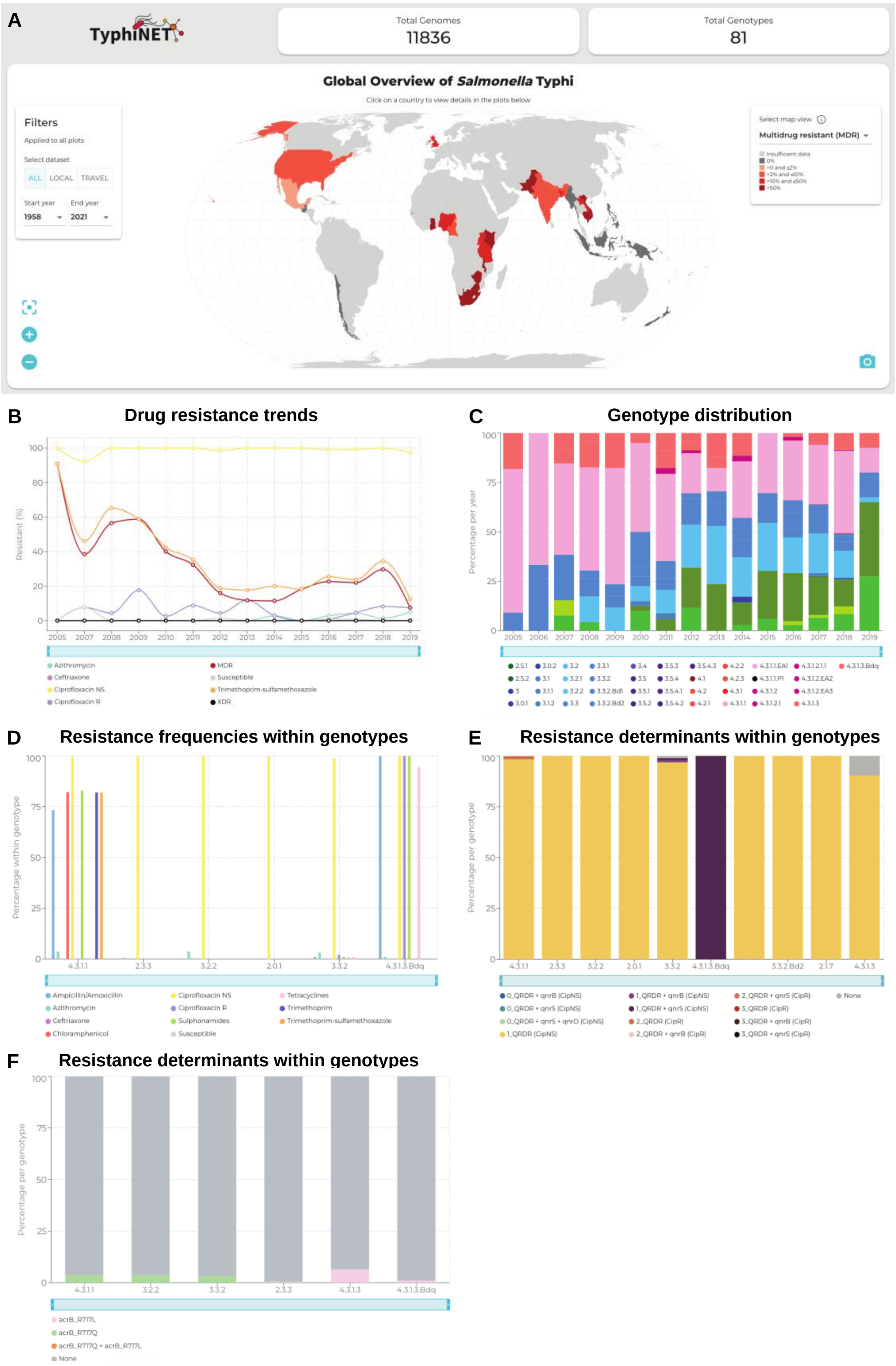
A
TyphiNET
Total Genomes
11836
Total Genotypes
81
Global Overview of Salmonella Typhi
Click on a country to view details in the plots below
Filters
Applied to all plots
Select dataset
ALL LOCAL TRAVEL
Start year 1958
End year 2021
Select map view
Multidrug resistant (MDR)
B
Drug resistance trends
Resistant (%)
Azithromycin
Ceftriaxone
Ciprofloxacin NS
Ciprofloxacin R
MDR
Susceptible
Trimethoprim-sulfamethoxazole
XDR
C
Genotype distribution
Percentage per year
D
Resistance frequencies within genotypes
Percentage within genotype
Ampicillin/Amoxicillin
Azithromycin
Ceftriaxone
Chloramphenicol
Ciprofloxacin NS
Ciprofloxacin R
Sulphonamides
Susceptible
Tetracyclines
Trimethoprim
Trimethoprim-sulfamethoxazole
E
Resistance determinants within genotypes
Percentage per genotype
0_QRDR + qnrB (CipNS)
0_QRDR + qnrS (CipNS)
0_QRDR + qnrS + qnrD (CipNS)
1_QRDR (CipNS)
1_QRDR + qnrB (CipNS)
1_QRDR + qnrS (CipNS)
2_QRDR (CipR)
2_QRDR + qnrB (CipR)
2_QRDR + qnrS (CipR)
3_QRDR (CipR)
3_QRDR + qnrB (CipR)
3_QRDR + qnrS (CipR)
None
F
Resistance determinants within genotypes
Percentage per genotype
acrB_R717L
acrB_R717Q
acrB_R717Q + acrB_R717L
None

**Figure S4. Decline of MDR and emergence of azithromycin resistance in Bangladesh. (A) Global map view of MDR typhoid.** Countries are coloured by the frequency of MDR cases as per the inset legend. **(B) Drug resistance trends over time plot shows frequencies of resistance to selected drug classes in Bangladesh per year.** Lines are coloured as per the inset legend. **(C) Annual genotype distribution plot shows the frequencies of pathogen genotypes circulating in Bangladesh per year.** Genotypes are coloured as per the inset legend. **(D) Resistance frequencies within genotypes plot shows frequencies of resistance to different drug classes, within common genotypes circulating in Bangladesh.** Bars are coloured according to the inset legend. **(E-F) Resistance determinants within genotypes plot shows the distribution of specific genes and mutations mediating resistance to a selected drug class, within common pathogen genotypes.** Bars are coloured as per the inset legend**.**

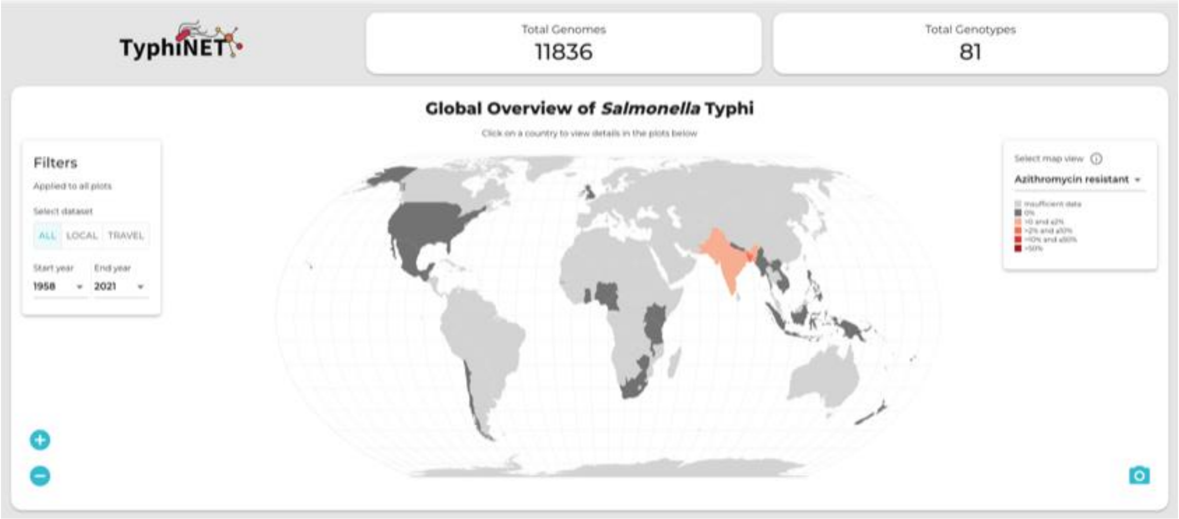


**Figure S5. Global map view of azithromycin resistant Typhi.** Countries on the map are coloured by azithromycin resistance frequency as per the inset legend (top right of map). Data are shown where there are ≥20 sequences available for the country of interest.

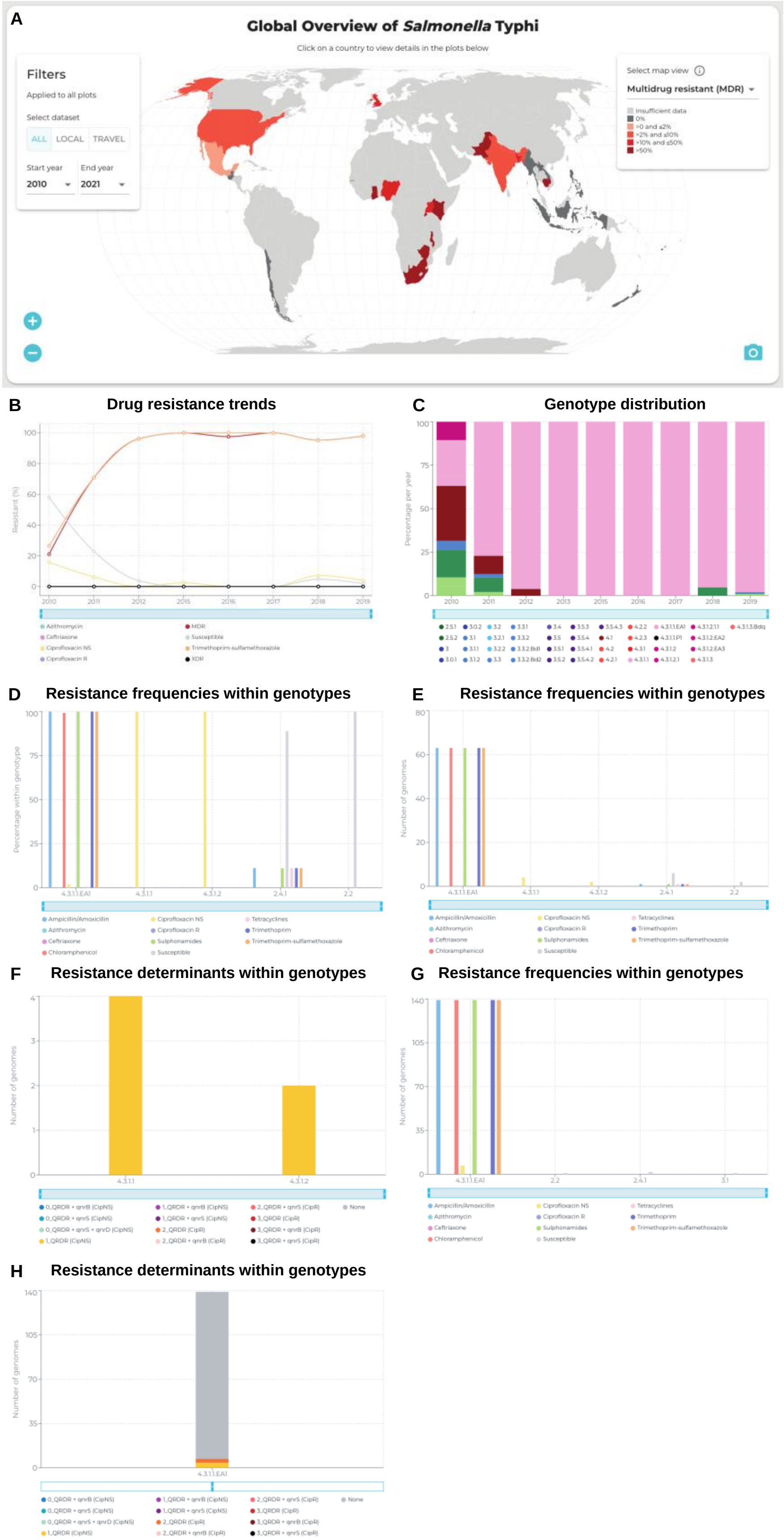
A
Global Overview of Salmonella Typhi
Click on a country to view details in the plots below
Filters
Applied to all plots
Select dataset
ALL
LOCAL
TRAVEL
Start year
2010
End year
2021
Select map view
Multidrug resistant (MDR)
B
Drug resistance trends
C
Genotype distribution
D
Resistance frequencies within genotypes
E
Resistance frequencies within genotypes
F
Resistance determinants within genotypes
G
Resistance frequencies within genotypes
H
Resistance determinants within genotypes

**Figure S6. Variation in genotype distribution and drug-resistance in Malawi from 2009. (A) Global map view of MDR typhoid from 2010 onwards.** Countries are coloured by the frequency of MDR cases as per the inset legend. **(B) Drug resistance trends over time plot shows frequencies of resistance to selected drug classes in Malawi per year.** Lines are coloured as per the inset legend. **(C) Annual genotype distribution plot shows the frequencies of pathogen genotypes circulating in Malawi per year.** Genotypes are coloured as per the inset legend. **(D) Resistance frequencies within genotypes plot shows frequencies of resistance to different drug classes, within common genotypes circulating in Malawi.** Bars are coloured according to the inset legend. **(E) Resistance frequencies within genotypes plot shows frequencies of resistance to different drug classes, within common genotypes circulating in Malawi between 2010 and 2012.** Bars are coloured as per the inset legend**. (F) Resistance determinants within genotypes plot shows the distribution of QRDR mutations mediating resistance to fluoroquinolones, within common pathogen genotypes between 2010 and 2012.** Bars are coloured as per the inset legend**. (G) Resistance frequencies within genotypes plot shows frequencies of resistance to different drug classes, within common genotypes circulating in Malawi between 2018 and 2019.** Bars are coloured as per the inset legend**. (H) Resistance determinants within genotypes plot shows the distribution of QRDR mutations mediating resistance to fluoroquinolones, within common pathogen genotypes between 2018 and 2019.** Bars are coloured as per the inset legend**.**
