## Supplementary Data 1 for "The TyphiNET data visualisation dashboard: Unlocking *Salmonella* Typhi genomics data to support public health"

### Global Overview of *Salmonella* Typhi

Fri May 24 2024 19:51

This report was generated at Fri May 24 2024 19:51, using TyphiNET (https://www.typhi.net), a data visualisation platform that draws genome-derived data on antimicrobial resistance and genotypes from Typhi Pathogenwatch (https://pathogen.watch), curated by the Global Typhoid Genomics Consortium (https://www.typhoidgenomics.org).

#### Source Data

TyphiNET data were last updated on March 24th 2024. For code and further details please see: (https://github.com/typhoidgenomics/TyphiNET).

TyphiNET presents data aggregated from >100 studies. Data for country Pakistan are drawn from studies with the following PubMed IDs (PMIDs) or Digital Object Identifier (DOI): 27069781, 25961941, 27703135, 32883020, 33704480, 33496224, 34370659, 31513580, 29216342, 34543095, 35750070.

Individual genome information, including derived genotype and AMR calls, sequence data accession numbers, and source information (PubMedID for citation) can be downloaded as a spreadsheet from the TyphiNET website (https://www.typhi.net).

#### Variable definitions

The genotypes reported here are defined in Dyson & Holt (2021), J. Infect. Dis. (https://doi.org/10.1093/infdis/jiab414).

Travel-associated cases are attributed to the country of travel, not the country of isolation, Ingle et al. 2019, PLoS NTDs., (https://doi.org/10.1371/journal.pntd.0007620).

Antimicrobial resistance determinants are described in the Typhi Pathogenwatch paper, Argimon et al. 2021, Nat. Commun., (https://doi.org/10.1038/s41467-021-23091-2).

#### Abbreviations

1. MDR, multi-drug resistant (resistant to ampicillin, chloramphenicol, and trimethoprim-sulfamethoxazole)

2. XDR, extensively drug resistant (MDR plus resistant to ciprofloxacin and ceftriaxone)

3. Ciprofloxacin NS, ciprofloxacin non-susceptible (MIC >=0.06 mg/L, due to presence of one or more *qnr* genes or mutations in *gyrA/parC/gyrB*)

4. Ciprofloxacin R, ciprofloxacin resistant (MIC >=0.5 mg/L, due to presence of multiple mutations and/or genes, see Carey et al, 2023 https://doi.org/10.7554/eLife.85867)

**Funding**

This project has received funding from the Wellcome Trust (Open Research Fund, 219692/Z/19/Z and AMRnet project, 226432/Z/22/Z) and the European Union Horizon 2020 research and innovation programme under the Marie Sklodowska-Curie grant agreement No 845681

### Global Overview of *Salmonella* Typhi

Total: 1526 genomes

Country: Pakistan

Time period: 1958 to 2021

---

**Map**

Map View: Ciprofloxacin non-susceptible (CipNS)

Dataset: All (local + travel)

Insufficient data
0%
>0 and ≤2%
>2% and ≤10%
>10% and ≤50%
>50%

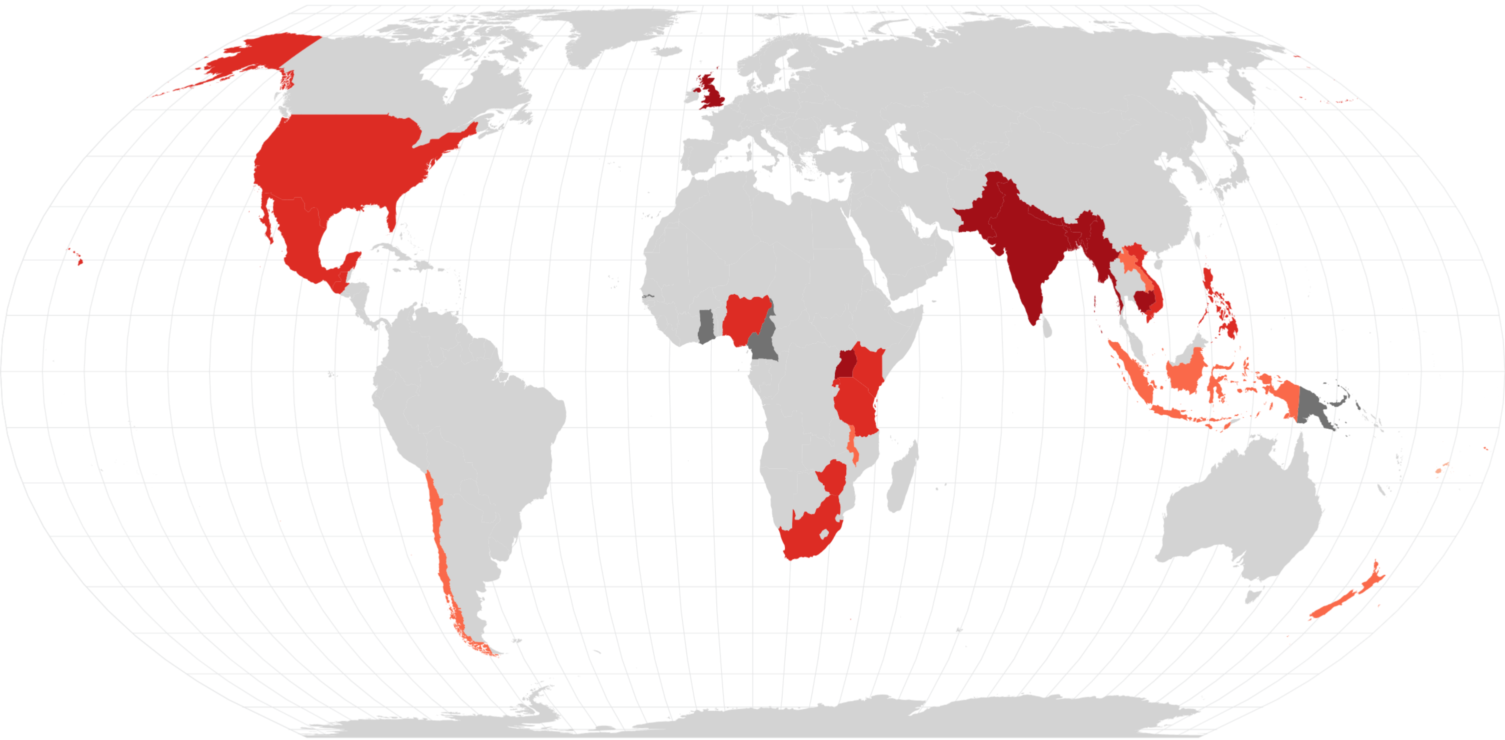

---

Source: typhi.net [Fri May 24 2024 19:51]

**Resistance frequencies within genotypes**

Top Genotypes (up to 7)

Total: 1526 genomes
Country: Pakistan
Time period: 1958 to 2021
Dataset: All (local + travel)

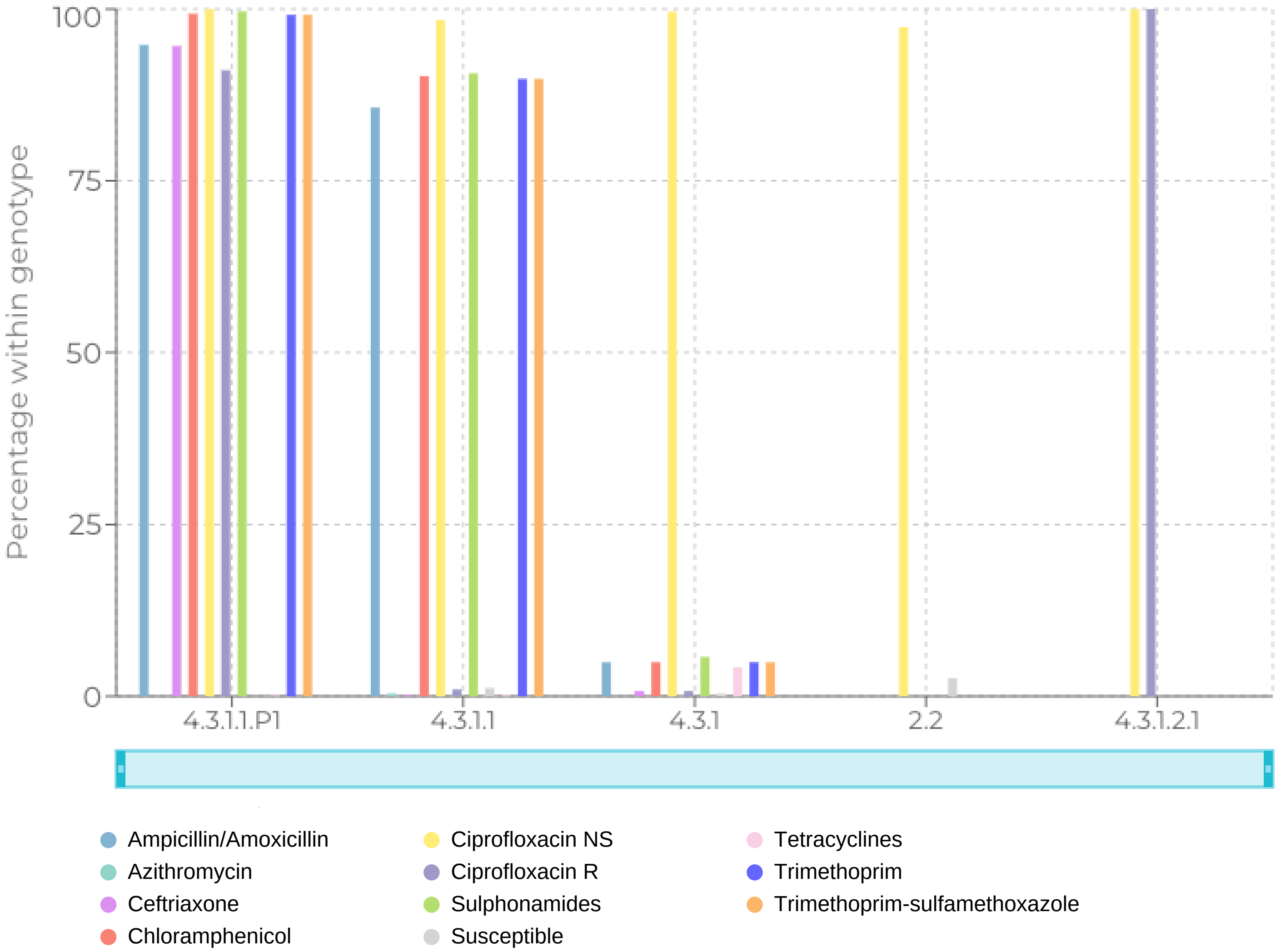

**Drug resistance trends**

Data are plotted for years with N >= 10 genomes

Total: 1526 genomes
Country: Pakistan
Time period: 1958 to 2021
Dataset: All (local + travel)

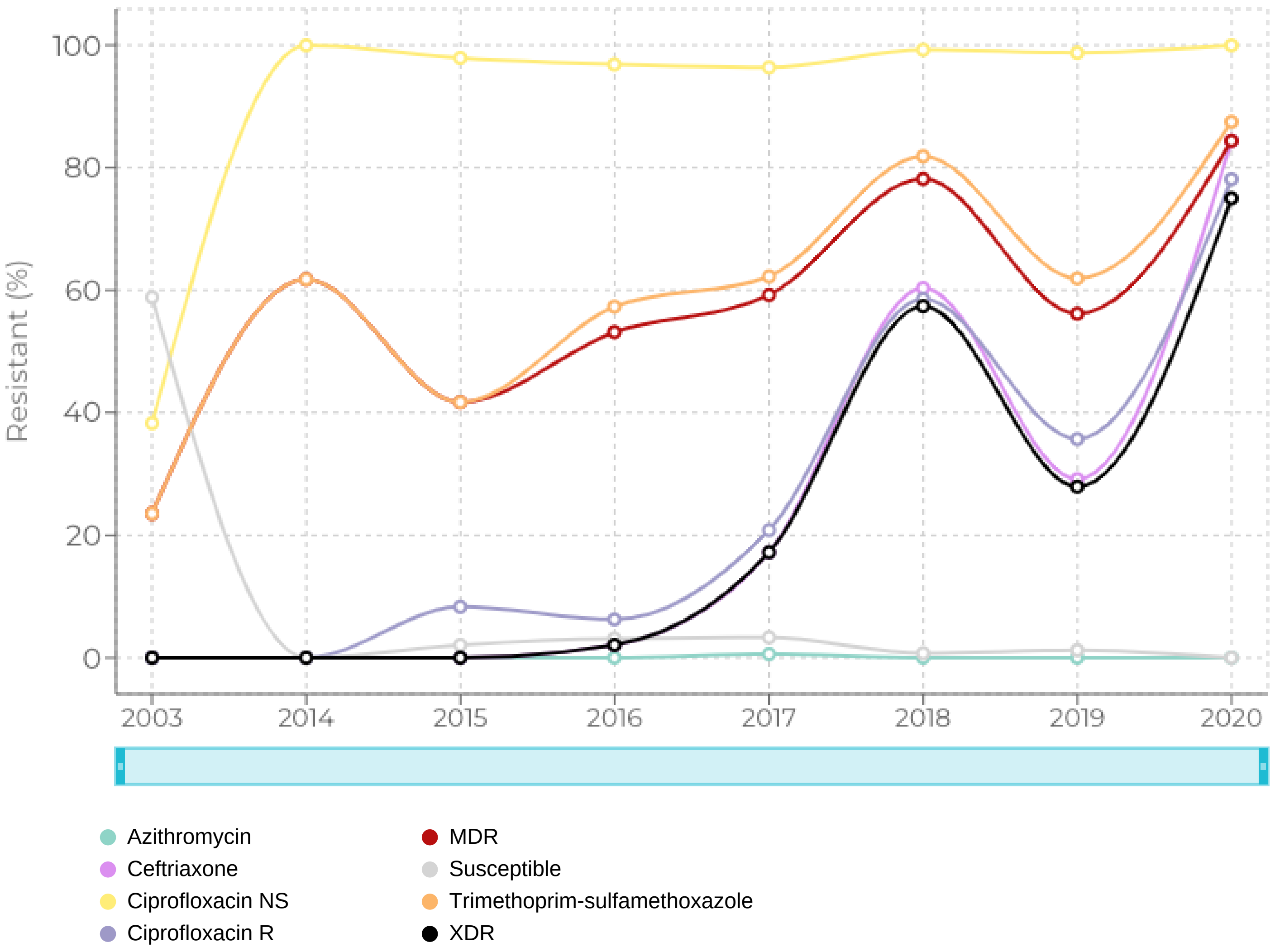

**Resistance determinants within genotypes: Ciprofloxacin NS**

Top Genotypes (up to 10)

Total: 1526 genomes
Country: Pakistan
Time period: 1958 to 2021
Dataset: All (local + travel)

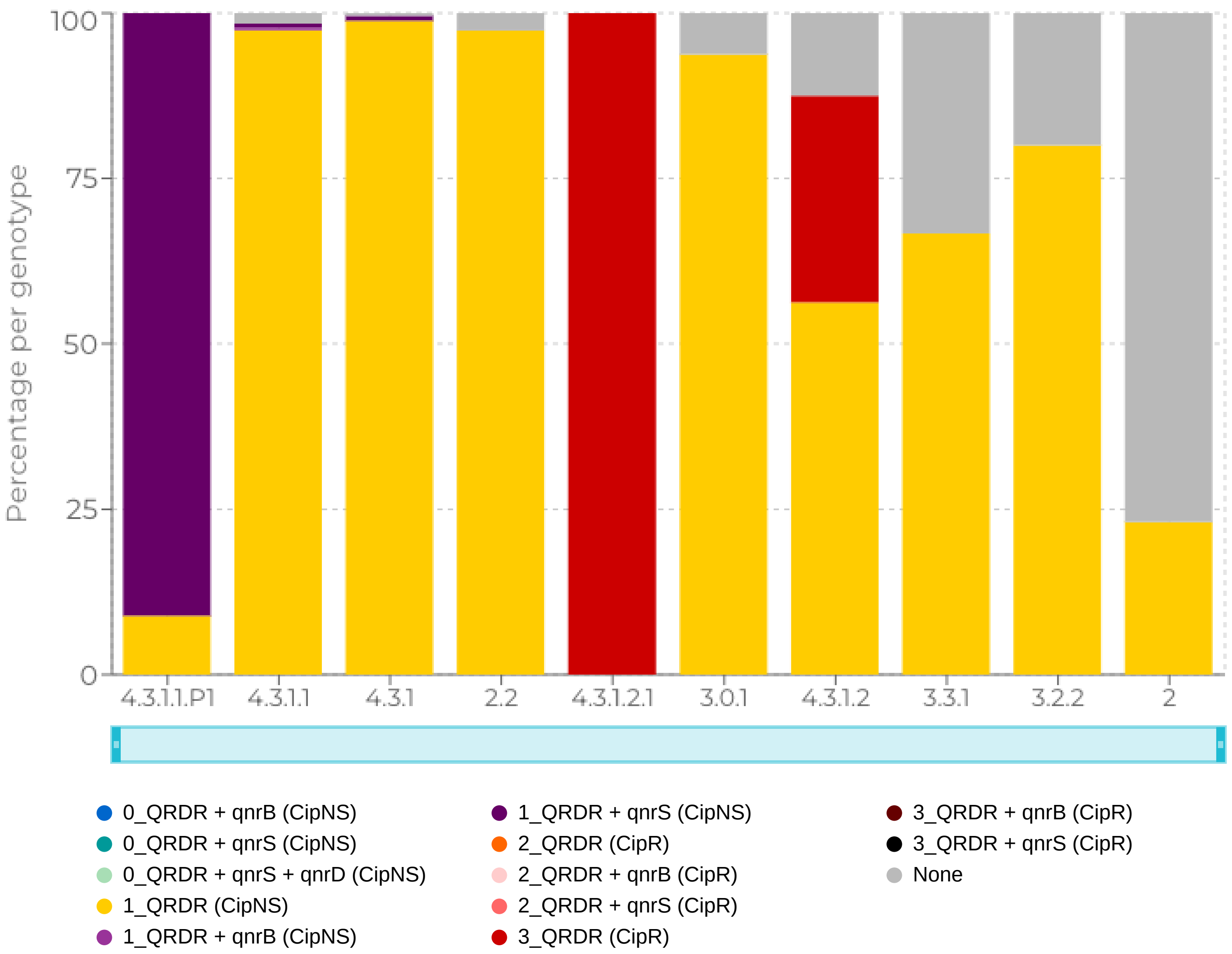

**Genotype distribution**

Total: 1526 genomes
Country: Pakistan
Time period: 1958 to 2021
Dataset: All (local + travel)

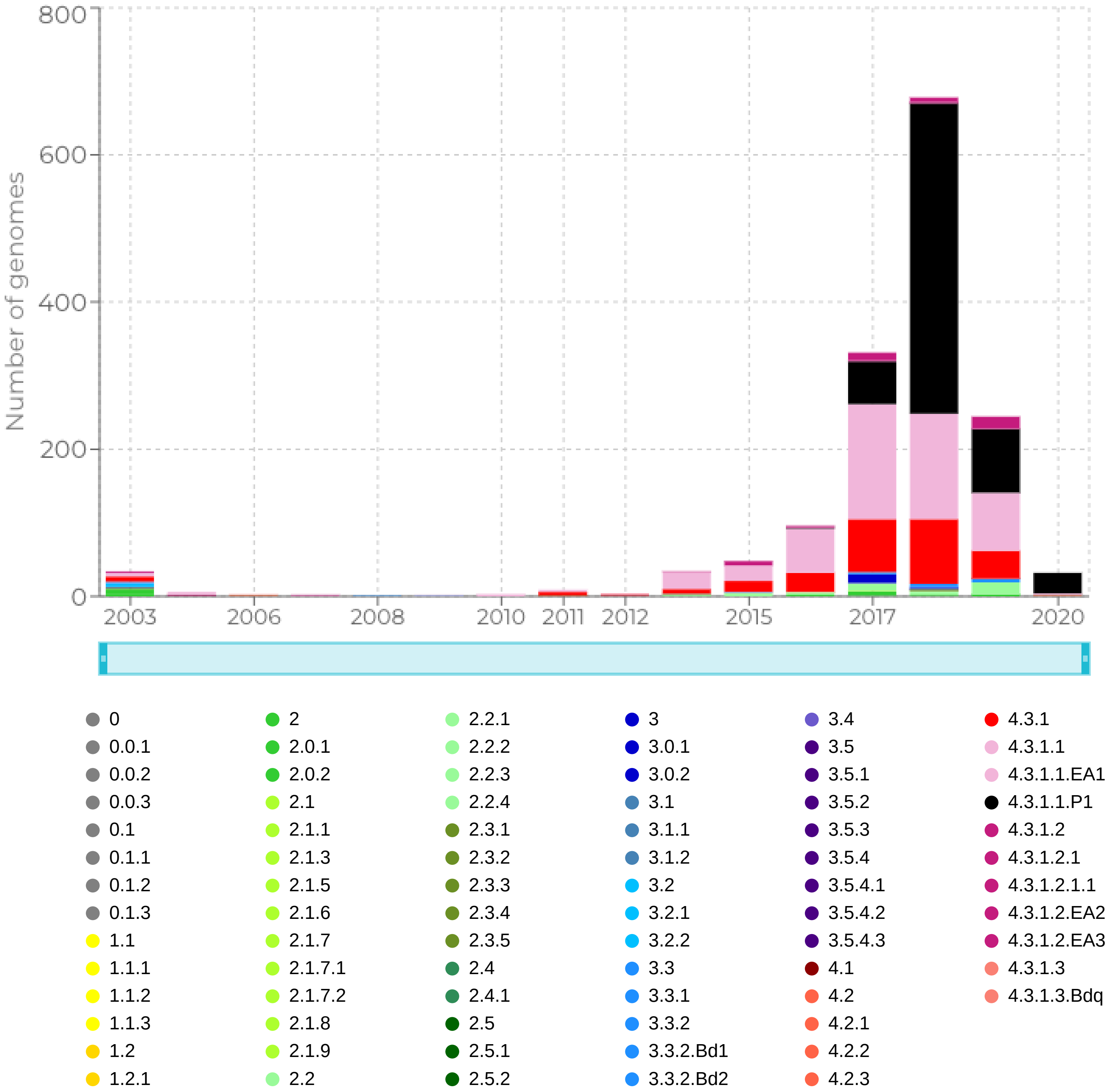
