## Supplementary figures and images for "The TyphiNET data visualisation dashboard: Unlocking *Salmonella* Typhi genomics data to support public health"

### Supplementary Figure 1

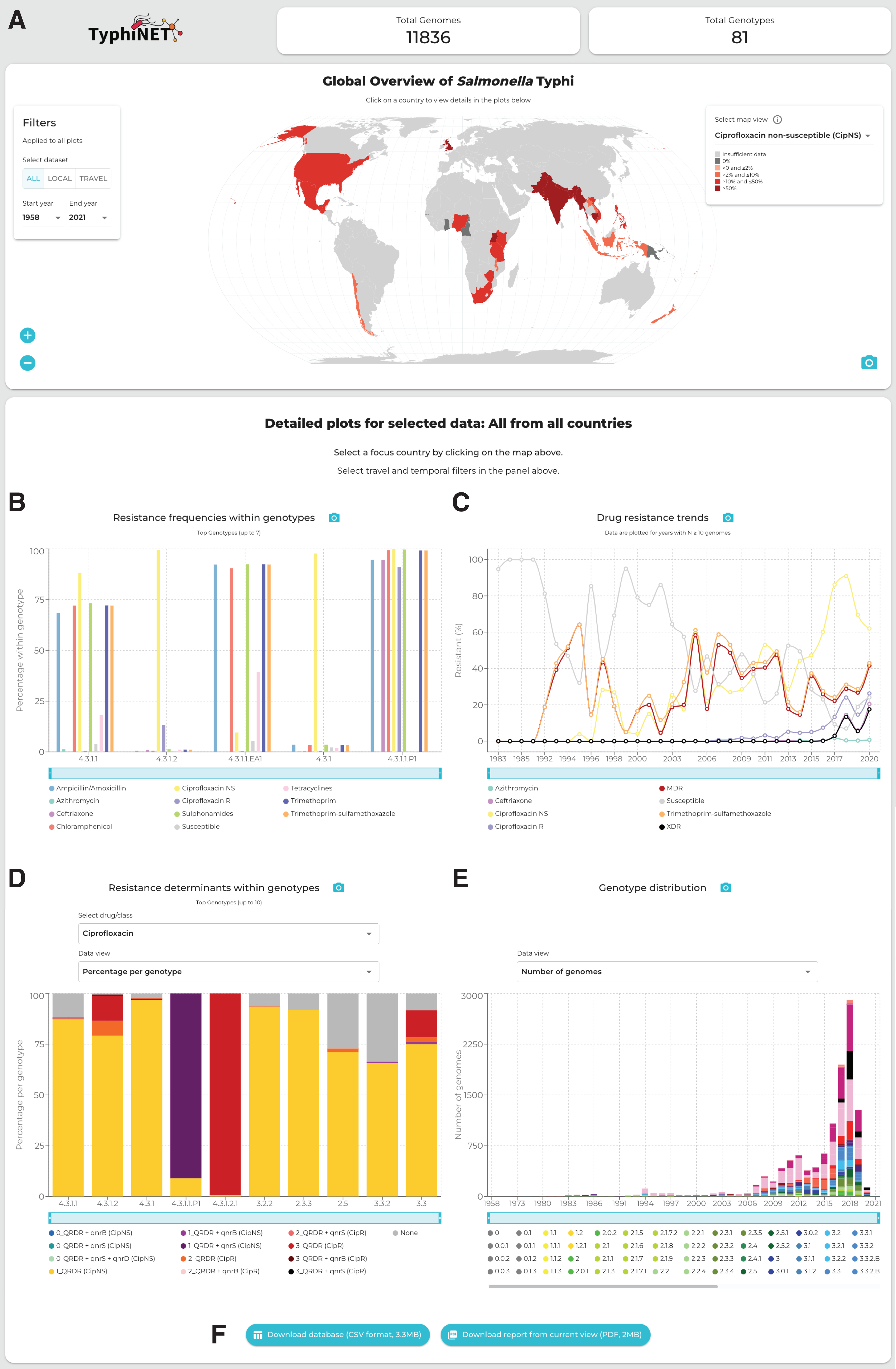

### Supplementary Figure 2

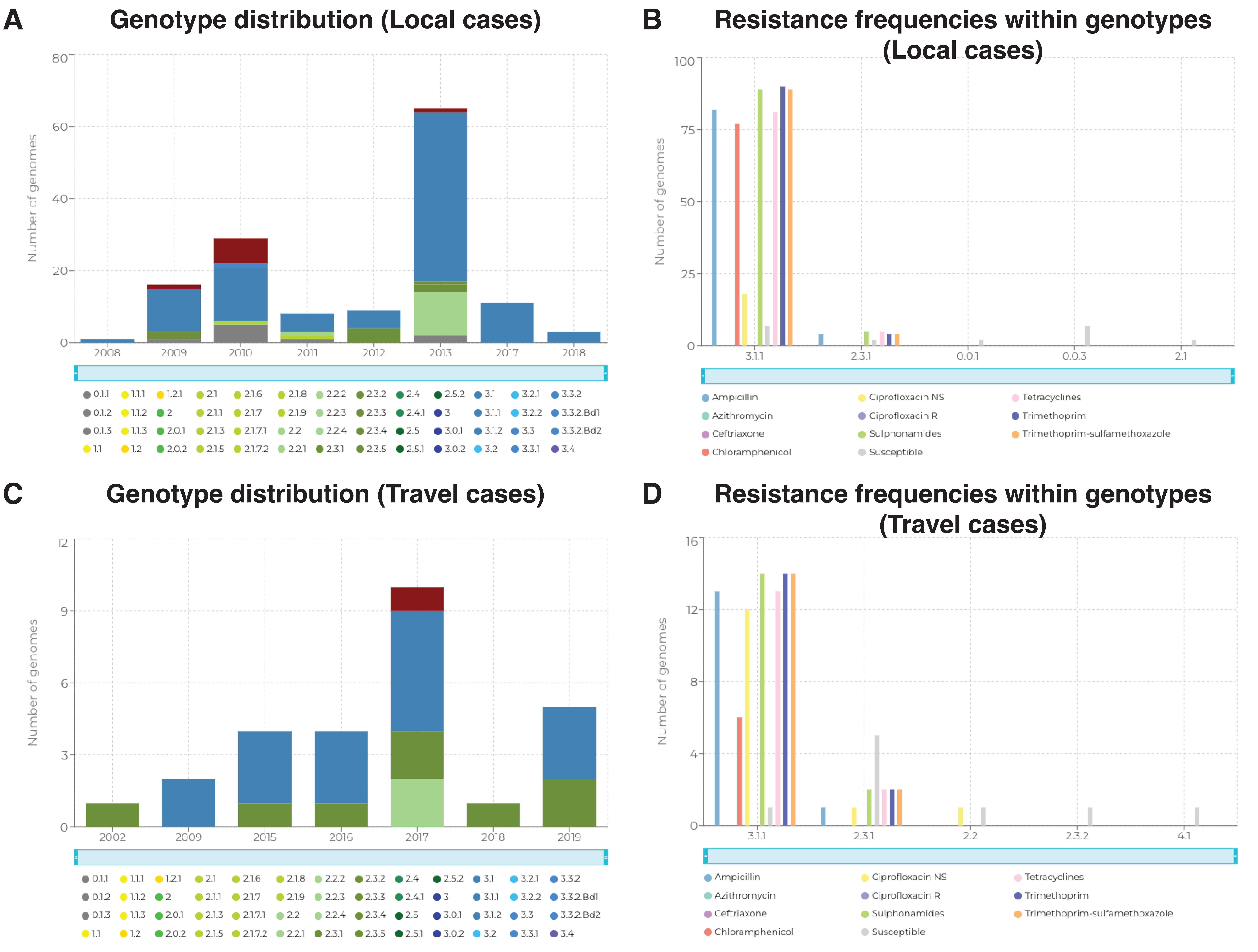

### Supplementary Figure 3

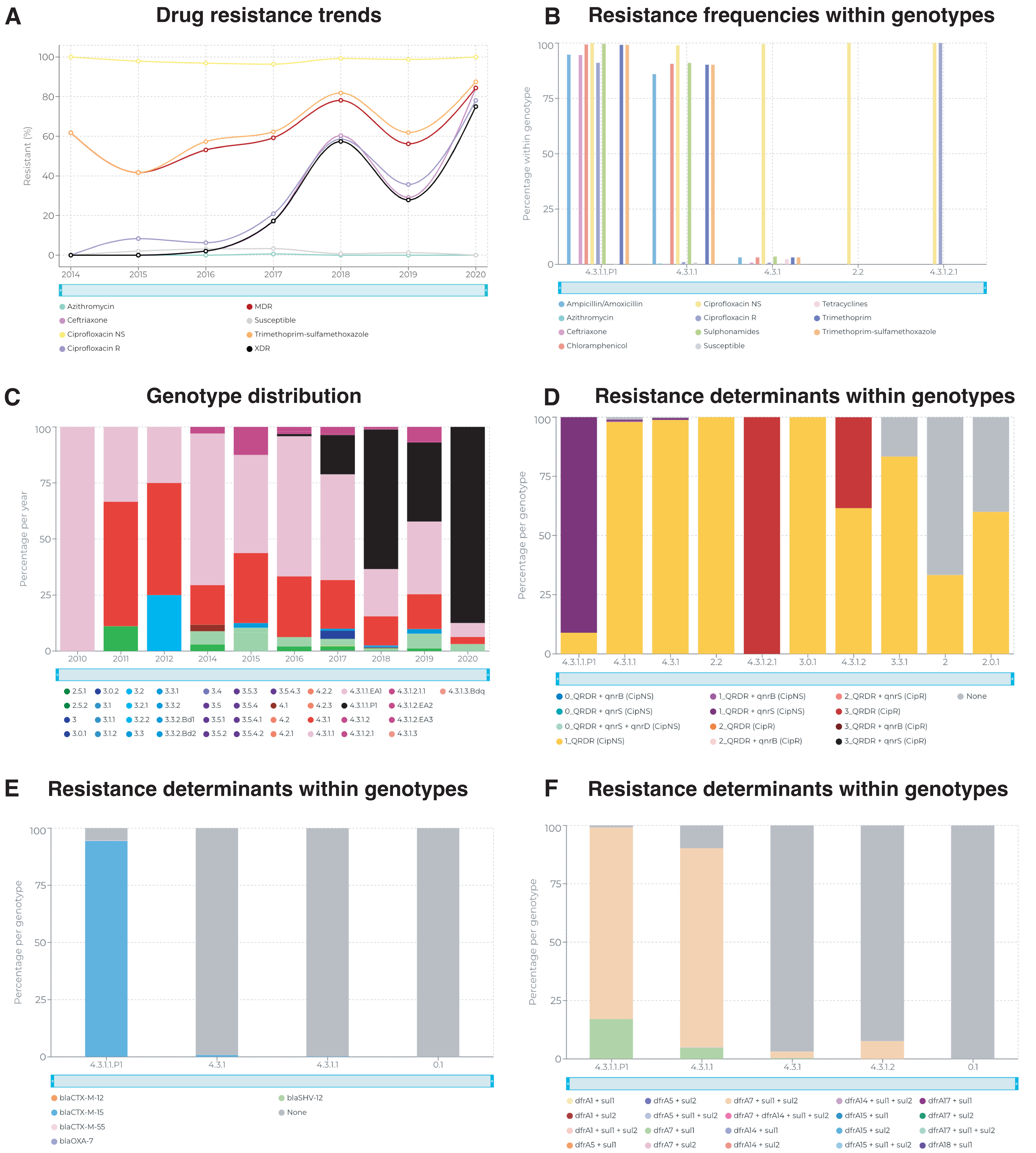

### Supplementary Figure 4

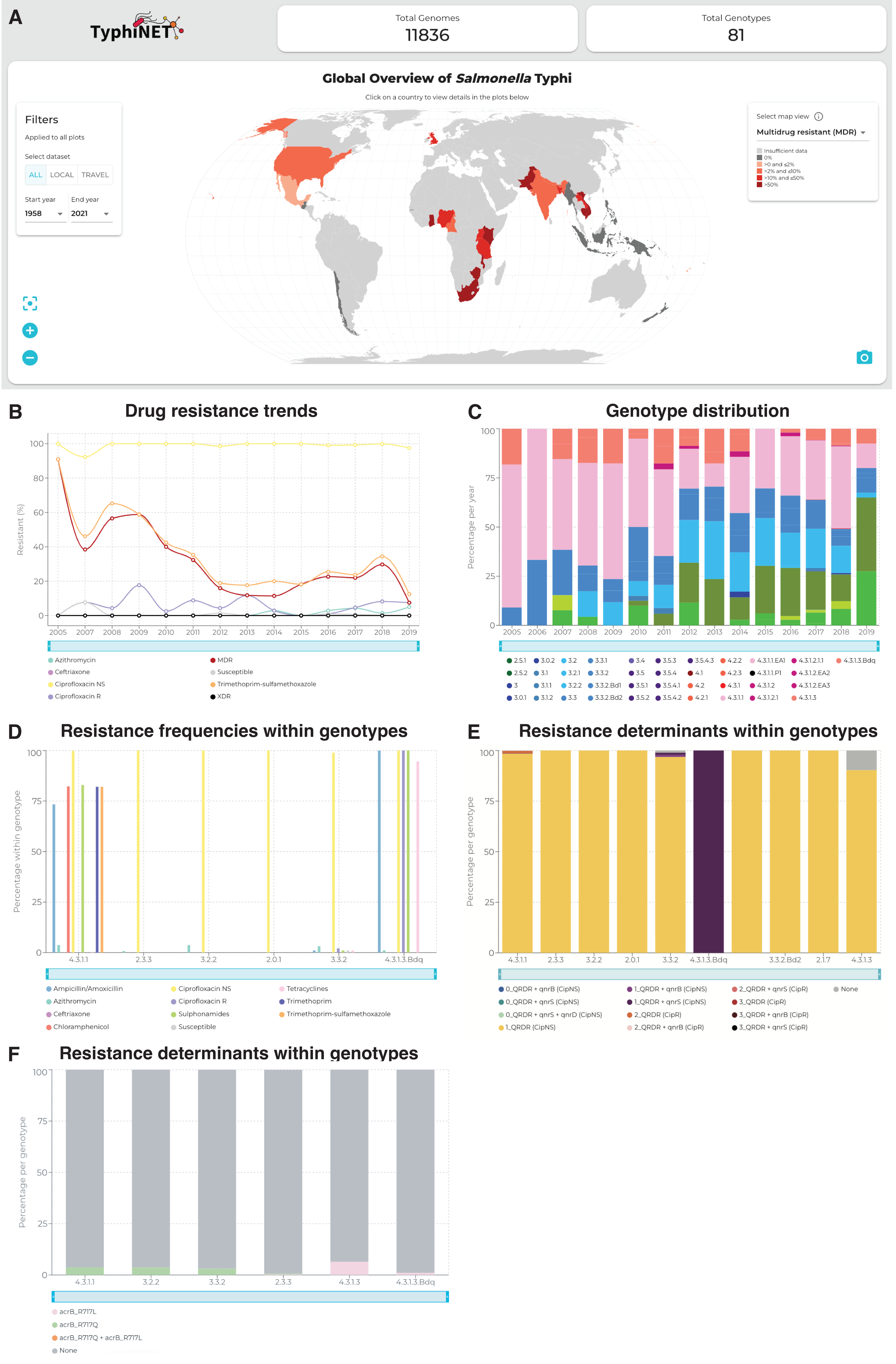

### Supplementary Figure 6

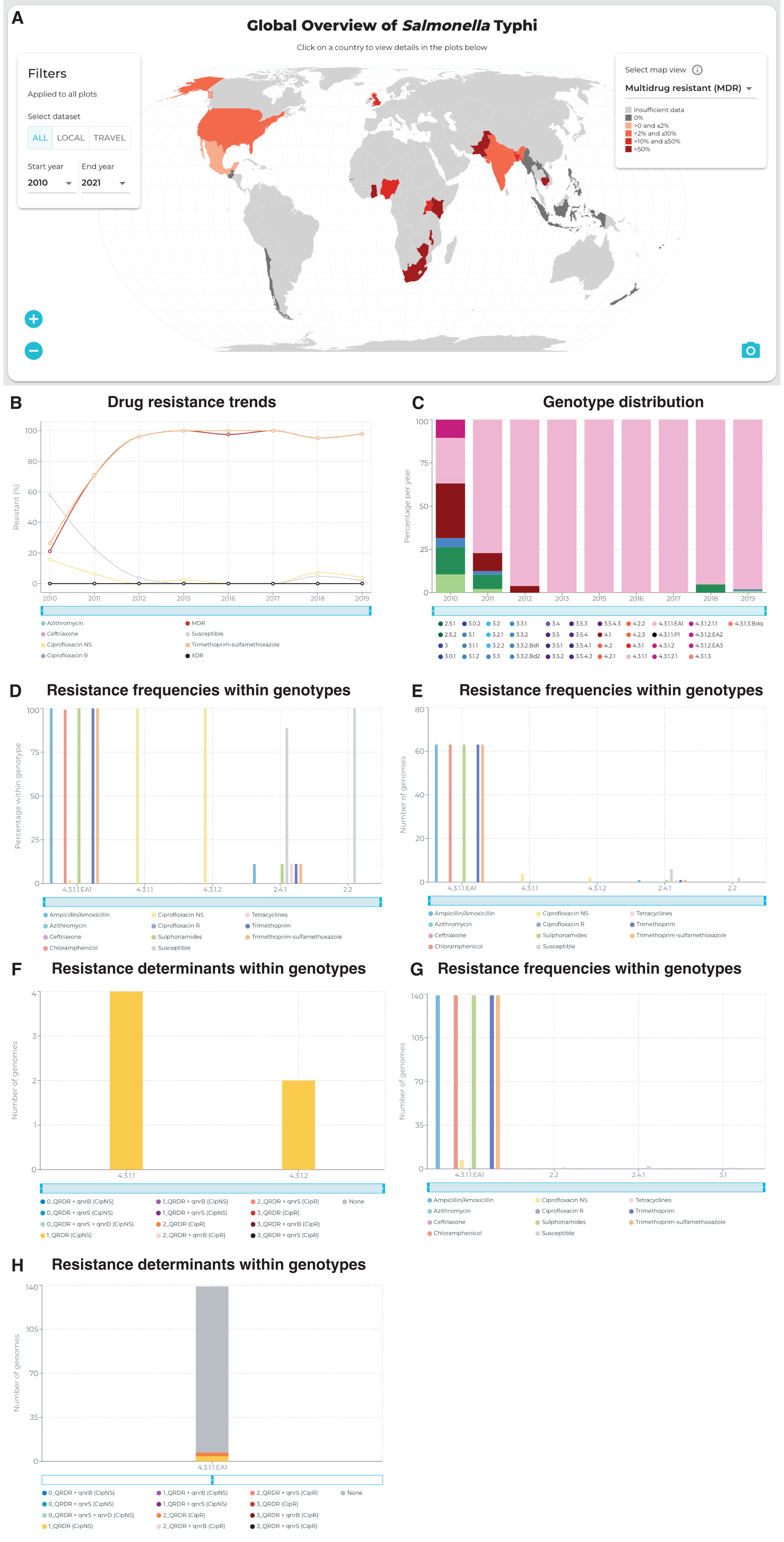
