## Supplementary Figure 5 for "The TyphiNET data visualisation dashboard: Unlocking *Salmonella* Typhi genomics data to support public health"

TyphiNET

Total Genomes
11836

Total Genotypes
81

### Global Overview of *Salmonella* Typhi

Click on a country to view details in the plots below

#### Filters

Applied to all plots

Select dataset

ALL | LOCAL | TRAVEL

Start year
1958

End year
2021

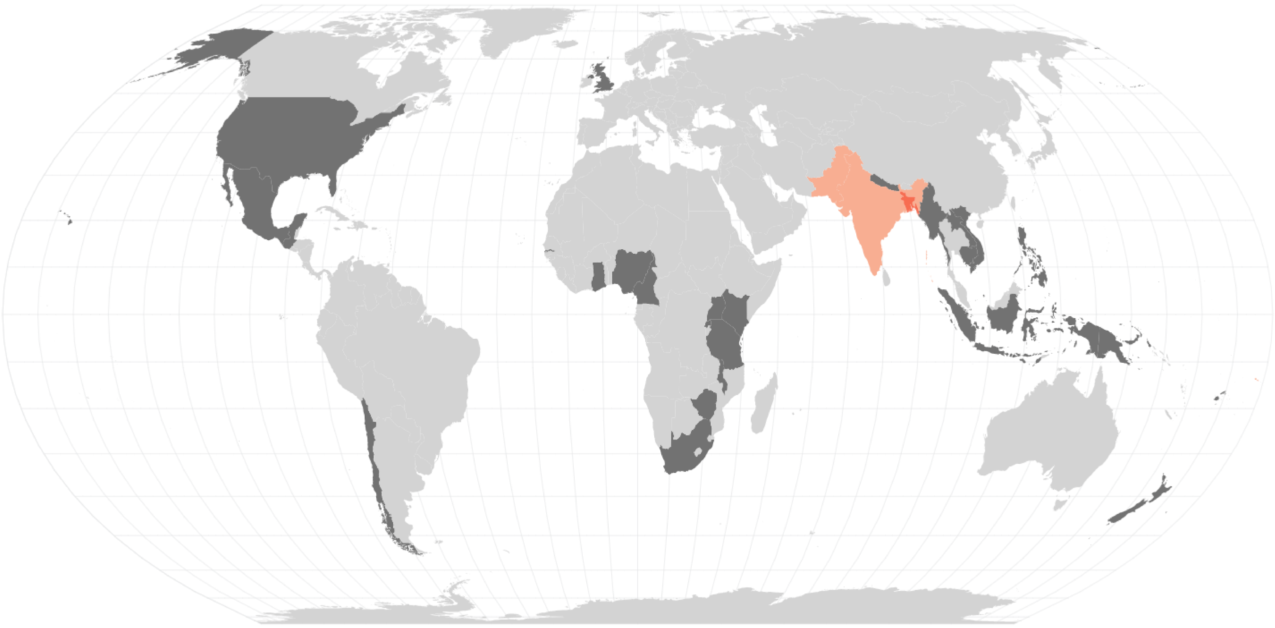

Select map view

Azithromycin resistant

Insufficient data
0%
>0 and ≤2%
>2% and ≤10%
>10% and ≤50%
>50%
